# TALAVE in Breast Cancer: *BRCA1/2* Mutation-Dependent Immune Remodeling after PARP Inhibition with Limited Checkpoint Engagement

**DOI:** 10.64898/2026.09.17.752386

**Authors:** Filipa Lynce, Kenichi Shimada, Claudine Isaacs, Xue Geng, Edward T. Richardson, Adam Nelson, Candace Mainor, Mei Wei, Julie M. Collins, Paula R. Pohlmann, Arielle L. Heeke, Kelly F. Zheng, Madeline G. Townsend, Nicole Swanson, Lauren M. Sloat, Jane Staunton, Stuart J. Schnitt, Hongkun Wang, Joan S. Brugge, Geoffrey I. Shapiro, Jennifer L. Guerriero

## Abstract

PARP inhibitors (PARPi) drive efficacy in BRCA-mutant breast cancer (BC) via DNA damage-induced synthetic lethality and immune activation, supporting their combination with immune checkpoint blockade. In the TALAVE study, patients with advanced BRCA-mutant or wild-type (WT) HER2-negative BC received talazoparib followed by talazoparib plus avelumab. Only BRCA-mutant BC responded clinically. Serial multi-omic profiling revealed BRCA-dependent tumor-immune remodeling, including sustained γH2AX–pTBK1 signaling with increased CD8+ T cells, tumor cell depletion, and enrichment of CD163+ macrophages. In contrast, BRCA-WT tumors remained compact and immunosuppressed with reduced T cells after therapy. PD-1+ T cells localized to CD4+-rich neighborhoods and correlated with longer progression-free survival in BRCA-mutant tumors. However, PD-L1+ cells were rapidly depleted or confined to immune-excluded regions, spatially segregating them from PD-1+ T cells, impairing effective checkpoint blockade. These findings suggest limited benefit of PD-1/PD-L1 blockade in augmenting PARPi activity and highlight the need for alternative strategies to sustain PARPi-induced immunity.

## Introduction

Genomic DNA is continually exposed to stress that causes damage from diverse endogenous and exogenous sources.^1^ Defects in DNA repair pathways allow lesions to progress to mutations and chromosomal abnormalities that increase cancer risk. *BRCA1* and *BRCA2* encode essential components of the homologous recombination (HR) pathway for repair of DNA double-strand breaks.^2^ Germline *BRCA1* and/or *BRCA2* (*gBRCA1/2*) mutations that cause functional loss increase lifetime risk of breast cancer to 50-80%^3–6^ from an average of 13% in the general population.^7^

BRCA-deficient breast cancers are highly sensitive to inhibitors of poly(ADP-ribose) polymerase (PARP), which prevent the repair of single-strand breaks, causing their collapse into double-strand breaks.^8, 9,10^ In *BRCA1/2*-mutant cells, which lack efficient HR repair, these breaks are lethal or are shunted to error-prone end-joining pathways, including microhomology-mediated end joining (MMEJ) and non-homologous end joining, resulting in marked chromosomal instability and ultimately cell death.^8, 11, 12^ PARP inhibition also traps PARP on DNA, forming toxic PARP-DNA complexes that cannot be bypassed in HR repair-deficient cells^13, 14^, and exacerbates single-strand DNA gap formation^15^, mechanisms also contributing to synthetic lethality.

The PARP inhibitors (PARPi) olaparib and talazoparib are approved by the U.S Food and Drug Administration (FDA) for patients with *gBRCA1/2*-mutated, HER2-negative metastatic breast cancer. Both agents significantly improve progression-free survival (PFS) compared with single-agent chemotherapy.^16, 17^ However, responses are rarely durable and have not translated into an overall survival (OS) advantage.^18, 19^ Acquired resistance emerges in nearly all patients, primarily through mechanisms of HR restoration mediated by BRCA reversion mutation or end resection rewiring. ^20–23^

The contribution of the breast tumor immune microenvironment to PARPi response and resistance remains incompletely understood. There is a growing body of evidence suggesting that BRCA genotype shapes baseline tumor immune activity.^24^ In a prior study, we showed that treatment-naïve *BRCA1*-mutant triple-negative breast cancers (TNBC) exhibit increased T cell and macrophage infiltration compared to *BRCA1/2*-wild-type tumors.^25^ This finding was consistent with endogenous DNA damage and activation of the DNA-sensing cGAS-STING signaling pathway reported in DNA repair-deficient primary breast cancers.^26^

In addition to these baseline differences between *BRCA1/2*-mutant and wild-type tumors, our preclinical studies in *BRCA1*-mutant human breast cancer cell lines and in a Brca1-deficient mouse model in which tumors were expanded in syngeneic immunocompetent mice, showed that exogenous DNA damage mediated by PARP inhibition induces cytosolic DNA accumulation further activating the cGAS-STING pathway, and promoting type I interferon-driven recruitment and activation of CD8+ T cells.^27^ While these findings are specific to BRCA-deficient cells in breast cancer models, similar results have been observed irrespective of BRCA deficiency in other cellular contexts.^28^ In Brca1-deficient mouse tumors, we further demonstrated that PARPi efficacy relies on both direct tumor-cell cytotoxicity and intratumoral cGAS-STING-mediated immune activation, with CD8+ T cell depletion markedly reducing treatment efficacy.^27^ In these tumors, PARP inhibition also results in infiltration of tumor-associated macrophages (TAMs) of complex phenotype, expressing immunosuppressive markers and utilizing lipid metabolic signaling. ^25^

It remains uncertain whether PARP inhibition routinely elicits similar immune modulation in human breast tumors, and whether such effects are sustained over time or translate to improved outcomes with the addition of immune checkpoint blockade. In MDA-MB-436 *BRCA1*-mutant xenografts expanded in humanized mice, the addition of PD-1 blockade to PARP inhibition resulted in only modestly improved efficacy.^29^ Additionally, early clinical trials combining PARPi with PD-1/PD-L1 inhibitors in metastatic breast cancer have yielded variable results.^30–33^ In the single-arm phase 2 TOPACIO/KEYNOTE-162, niraparib plus pembrolizumab showed modest activity in metastatic TNBC; responses were more frequent in *BRCA1/2-*mutated tumors, but also occurred in WT tumors.^30^ In the phase 1/2 MEDIOLA trial, olaparib plus durvalumab demonstrated encouraging activity in *gBRCA1/2*-mutated HER2-negative metastatic breast cancer, but the lack of a control arm limited assessment of the specific contribution of immune checkpoint blockade.^31^ Nonetheless, in another study of olaparib and durvalumab employing serial biopsies, destruction of a *BRCA1/2*-mutant basal breast cancer in a long-term survivor was accompanied by CD8+ T cell infiltration.^34^ By contrast, preliminary reported results of the randomized NCI-sponsored study of olaparib with or without atezolizumab in *BRCA1/2*-mutated advanced breast cancer (CTEP 10020) did not show improved PFS or OS with the addition of PD-L1 blockade across the population or in subsets with hormone receptor-positive or triple-negative BRCA-associated BC.^33^

Collectively, these data highlight the need to better define which tumors undergo meaningful immune remodeling after PARPi exposure with comprehensive characterization of the effects of PARP inhibition on the breast tumor microenvironment (TME) to inform the development of optimal immunologic combination strategies. To address these questions, we conducted an open-label, multi-institutional pilot study (TALAVE) evaluating the safety, efficacy, and immunologic effects of combining the PARPi talazoparib with the anti-PD-L1 antibody avelumab in patients with *BRCA1/2*-mutant (BRCA-MUT) BC or *BRCA1/2*-wild-type (BRCA-WT) HER2-negative advanced triple-negative breast cancer (TNBC). The initial rationale for inclusion of the BRCA-WT TNBC arm was to determine whether PARPi-based induction and combination with immunotherapy could stimulate immune responses in a BC subset less responsive to single-agent PARP inhibition.

Longitudinal assessment of the TME pre-treatment, post-talazoparib, and post-combination therapy demonstrated PARPi-mediated immune modulation in BRCA-MUT disease, consistent with preclinical models. However, PD-L1+ cells were depleted or confined to immune-excluded regions, resulting in spatial segregation from PD-1+ T cells and impairment of effective checkpoint blockade. These results may at least in part explain why the addition of standard immune checkpoint blockade has not consistently enhanced the clinical benefit of PARP inhibition in BRCA-associated breast cancer.

## Results

### Talazoparib plus avelumab is highly effective in BRCA-MUT breast cancer, with modest activity in BRCA-WT TNBC

#### Patient Population

This multi-institutional phase 1/2 trial enrolled patients with advanced HER2-negative breast cancer. The study was registered on ClinicalTrials.gov on April 22, 2019. The first patient provided informed consent on April 19, 2019, and initiated study treatment on April 30, 2019. The data cutoff for this analysis was June 2024. Baseline patient characteristics are summarized in **Table 1**. Between May 2019 and September 2022, 24 patients were enrolled, with 12 assigned to each cohort: Cohort 1 (BRCA-MUT BC) and Cohort 2 (BRCA-WT, TNBC) (**Fig. 1A**). The median age was 50 years (range: 27 – 78 years), and all patients were female. Participants had received a median of one prior line of therapy for advanced disease (range: 0 – 7), and 42% had previously received platinum-based chemotherapy. In Cohort 1, five patients had a deleterious germline *BRCA1* mutation, six had a germline *BRCA2* mutation, and one patient had tumor harboring somatic *BRCA2* mutation. Eight (66.7%) patients had hormone receptor-positive/HER2-negative BC and four patients (33.3%) had TNBC. All patients in Cohort 2 (100%) had TNBC (**Supplementary Table 1**).

**Figure 1.**
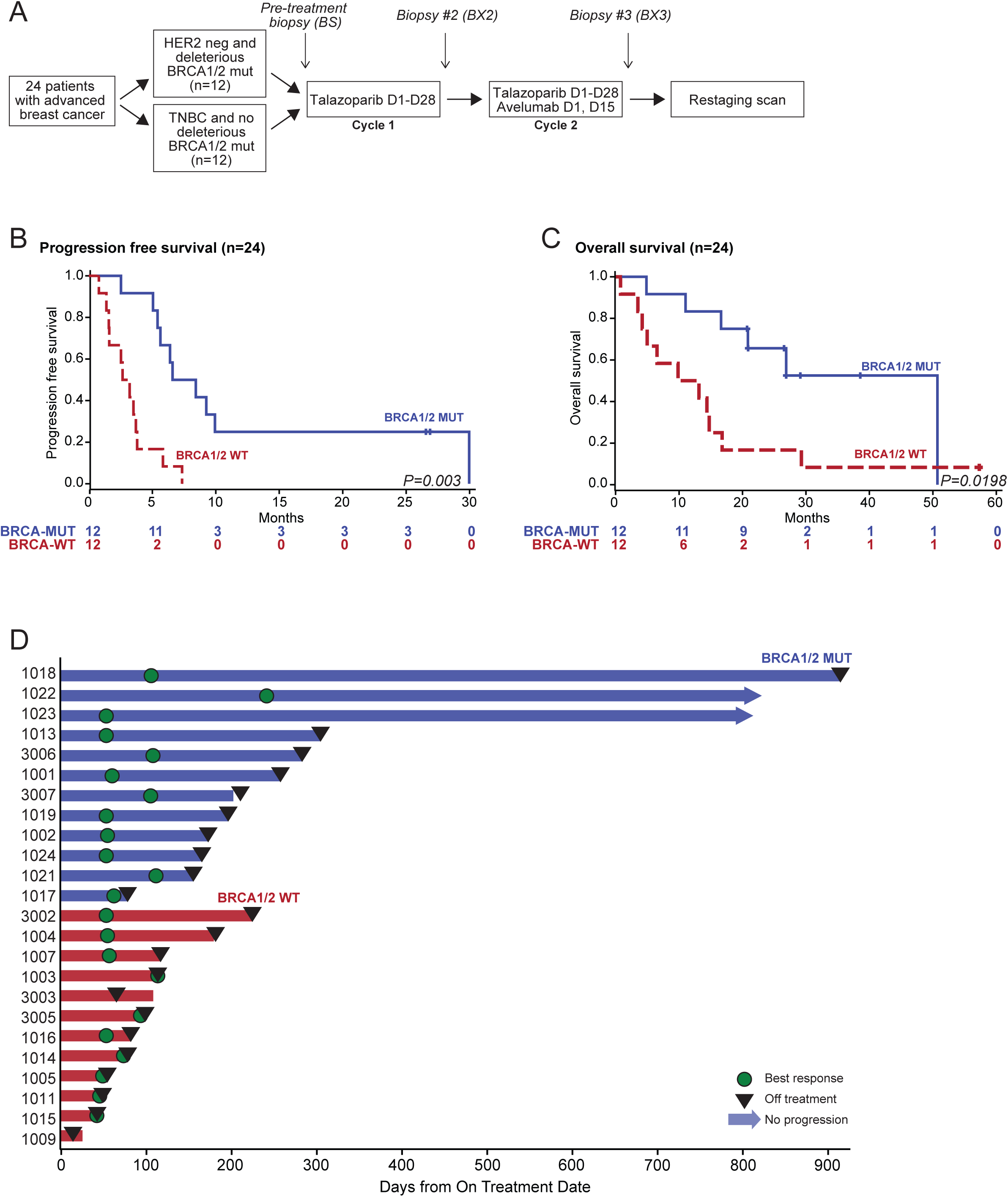
Clinical outcomes of PARP inhibitor therapy in BRCA-MUT tumors. (A) Schematic of the TALAVE trial design. (B) Kaplan–Meier curve for PFS, stratified by BRCA status. (C) Kaplan–Meier curve for OS, stratified by BRCA status. (D) Swimmer plot showing individual PFS durations for all 24 patients. P values are from two-sided log-rank tests comparing PFS or OS between the indicated groups.

**Table 1.**
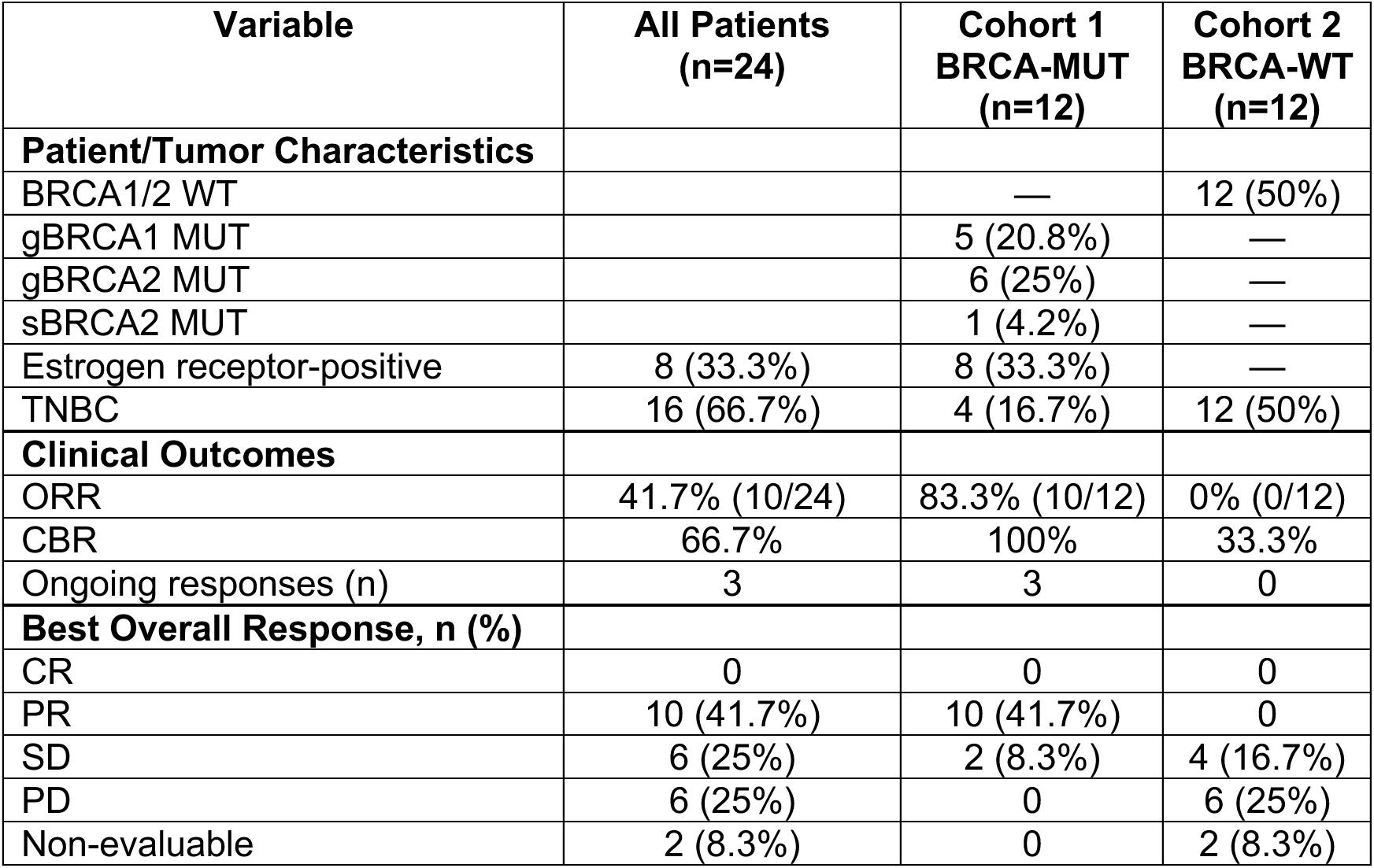
Patient/tumor characteristics and clinical outcomes by cohort.

| Variable | All Patients<br>(n=24) | Cohort 1<br>BRCA-MUT<br>(n=12) | Cohort 2<br>BRCA-WT<br>(n=12) |
| --- | --- | --- | --- |
| **Patient/Tumor Characteristics** |  |  |  |
| BRCA1/2 WT |  | — | 12 (50%) |
| gBRCA1 MUT |  | 5 (20.8%) | — |
| gBRCA2 MUT |  | 6 (25%) | — |
| sBRCA2 MUT |  | 1 (4.2%) | — |
| Estrogen receptor-positive | 8 (33.3%) | 8 (33.3%) | — |
| TNBC | 16 (66.7%) | 4 (16.7%) | 12 (50%) |
| **Clinical Outcomes** |  |  |  |
| ORR | 41.7% (10/24) | 83.3% (10/12) | 0% (0/12) |
| CBR | 66.7% | 100% | 33.3% |
| Ongoing responses (n) | 3 | 3 | 0 |
| **Best Overall Response, n (%)** |  |  |  |
| CR | 0 | 0 | 0 |
| PR | 10 (41.7%) | 10 (41.7%) | 0 |
| SD | 6 (25%) | 2 (8.3%) | 4 (16.7%) |
| PD | 6 (25%) | 0 | 6 (25%) |
| Non-evaluable | 2 (8.3%) | 0 | 2 (8.3%) |

#### Trial Design

Patients received talazoparib (1mg by orally [PO] once daily on days 1-28) as monotherapy during cycle 1, followed by combination therapy beginning in cycle 2 and continuing in all subsequent cycles. Avelumab (800 mg intravenously [IV] on day 1 every 2 weeks) was added to talazoparib (**Fig. 1A**). The phase 1 portion evaluated a single dose level, using the recommended phase 2 dose of each agent previously established in single-agent studies in patients with repair-proficient, *BRCA1/2*-mutant and *ATM*-mutant tumors.^19, 35, 36^ All six patients enrolled in the phase 1 portion were monitored for dose-limiting toxicities (DLT) during the first two cycles, as specified in the protocol. No DLTs were observed, and accrual subsequently proceeded to the phase 2 portion in accordance with the protocol.

#### Safety and Tolerability

Treatment-related adverse events (TRAEs) are listed for the overall population and by cohort in **Supplementary Table 2**. The most common all-grade TRAEs in the overall study population included nausea (37.5%), fatigue (37.5%), anemia (33.3%), and neutropenia (25%). The most common grade ≥ 3 TRAEs included anemia (29.2%), neutropenia (12.5%), and thrombocytopenia (12.5%).

#### Efficacy

In the overall study population, the objective response rate (ORR) was 41.7%, the clinical benefit rate (CBR; CR or PR, or SD lasting ≥ 24 weeks) was 66.7% and the median duration of response (DoR) was 5.2 months (95% CI 3.7-NE). Among patients in Cohort 1, 10 experienced a partial response (PR), for an ORR of 83.3%. An additional two patients in Cohort 1 experienced stable disease (SD) as the best response, for a CBR of 100% for the cohort. Among patients in Cohort 2, four experienced SD as the best response, for an ORR of 0% and a CBR of 33.3%. At the time the data were analyzed, there were two ongoing responders in Cohort 1 (16.7%; **Table 1**).

In the overall study population, the median PFS was 5.2 [95% CI: 2.7 – 6.6] months (**Supplementary Fig. 1A**). When stratified by BRCA status, patients in Cohort 1 (BRCA-MUT) experienced significantly longer PFS than those in Cohort 2 (BRCA-WT), with median PFS of 7.5 [95% CI: 5.1 – not estimable (NE)] versus 2.9 [95% CI: 1.4 – 3.8] months, respectively (**Fig. 1B**), consistent with prior clinical experience showing greater PARP inhibitor activity in BRCA-mutated, homologous recombination repair-deficient tumors. Among patients with *BRCA1/2* mutations, those with hormone receptor-positive disease had longer PFS compared to those with TNBC (**Supplementary Fig. 1B**). The median OS in the overall population was 16.7 [95% CI: 9.8 – 29.3] months (**Supplementary Fig. 1C**). When stratified by BRCA status, the median OS was 50.7 [95% CI: 11.0 – NE] months in Cohort 1 compared with 11.5 [95% CI: 3.5 – 16.8] months in Cohort 2 (**Fig. 1C**). Although the association between BRCA status and OS was less pronounced than that observed for PFS, it remained significant (**Fig. 1C, Supplementary Fig. 1D**). Three patients, all in cohort 1 (patients 1018, 1022, and 1023) with hormone receptor-positive disease, remained on study for more than two years (**Fig 1D**).

### Serial multi-omic profiling to identify correlatives of response to PARPi and PD-L1 blockade

To define how PARP inhibition and PD-L1 blockade remodel tumor and immune responses in patients, we analyzed tumor samples collected at baseline (BS), after one month of talazoparib monotherapy (BX2), and following one additional month of talazoparib plus avelumab combination therapy (BX3). Transcriptional profiling was performed using the PanCancer Immuno-Oncology 360^TM^ (IO360) panel, assessing 750 genes supplemented with 55 DNA damage response (DDR)-related genes (**Supplementary Data 1**). Spatial protein profiling was conducted using the Digital Spatial Profiling (DSP) Protein assay (52 proteins), and single-cell cyclic immunofluorescence (CyCIF) assessing 18 protein markers (**Fig. 2A**). Overall, samples from 8 BRCA-MUT and 9 BRCA-WT patients were analyzed, with each patient contributing to at least one dataset at one time point (**Supplementary Fig. 2A**). All available samples meeting quality control criteria were included in the analysis, irrespective of cross-platform overlap, ensuring comprehensive representation across modalities.

**Figure 2.**
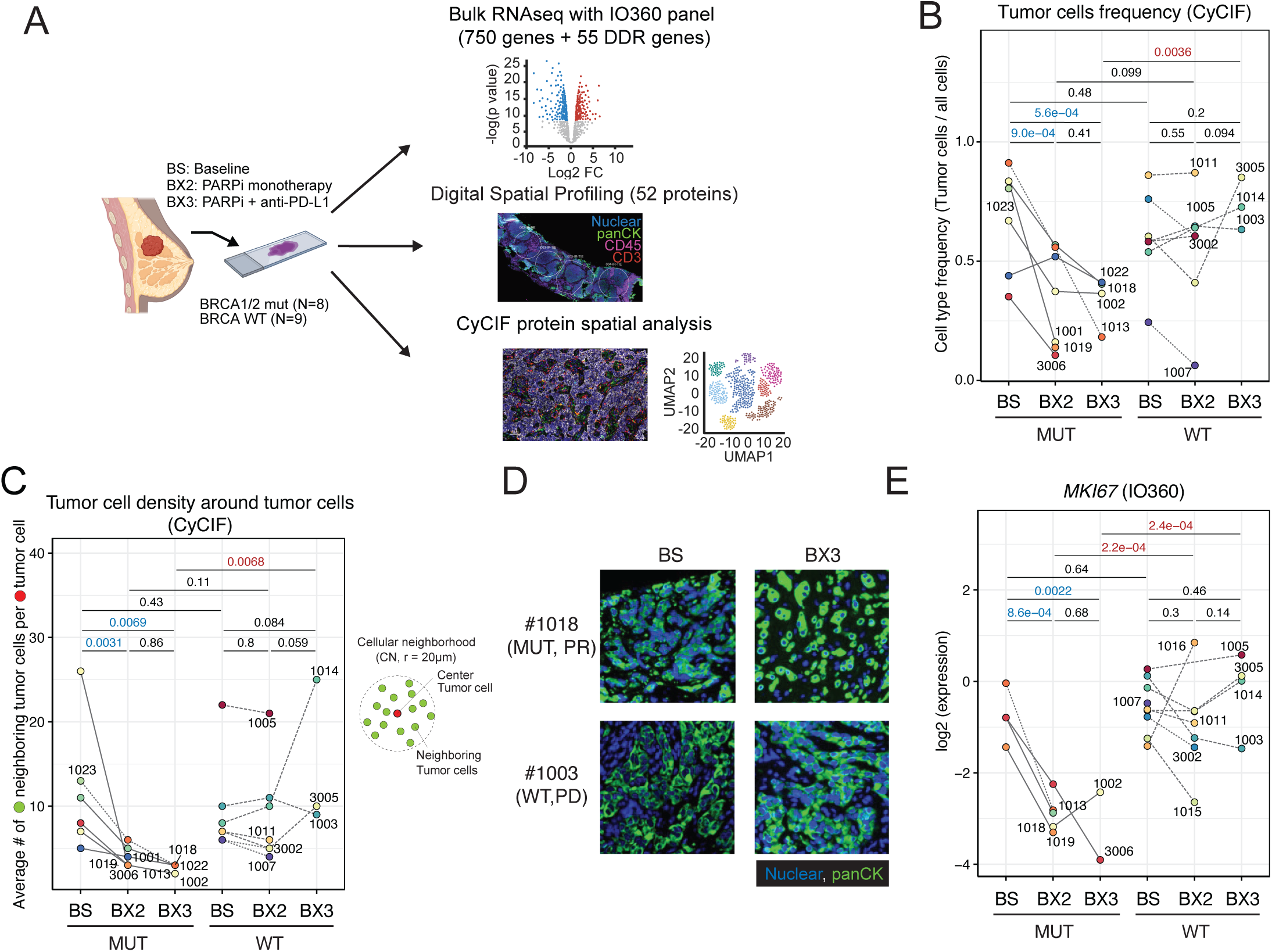
PARP inhibition depletes tumor cells and disrupts tumor architecture in BRCA-MUT but not BRCA-WT tumors. (A) Overview of sample collection and multimodal analysis workflow. (B) Tumor cell frequency as a proportion of all cells, stratified by BRCA status and treatment time point. Lines connect longitudinal samples from the same patient. (C) Tumor cell density within local cellular neighborhoods (CN; average number of tumor cells within a 20 μm fixed-radius surrounding each tumor cell in a sample). (D) Representative images of panCK (green) staining showing tumor architecture in BRCA-MUT and BRCA-WT patients at baseline and after combination therapy. (E) MKI67 expression in IO360. P values in B, C, and E are from two-sided linear mixed-effects models with patient as a random effect.

### Tumor architecture is disrupted in BRCA-MUT but remains intact in BRCA-WT tumors

To assess treatment-induced remodeling of the TME, we first analyzed cell composition and spatial organization of tumor cells across BRCA subgroups using CyCIF imaging. Computationally segmented cells were classified into major lineages based on marker expression using a predefined framework (**Supplementary Fig. 2B–D**). Longitudinal analysis of cell type frequencies in biopsies across time points using a linear mixed-effects regression model (LMER) revealed a significant decrease in tumor cell frequency in all BRCA-MUT tumors, whereas most BRCA-WT tumors showed minimal change in response to treatment (**Fig. 2B**). To determine whether this decline reflected structural disruption, we quantified the local tumor cell density by counting neighboring tumor cells within a 20 μm cellular neighborhood (CN) centered on each tumor cell. Mean tumor cell CN density was significantly reduced in BRCA-MUT tumors but remained stable or slightly increased in BRCA-WT tumors (**Fig. 2C,D**).

We confirmed these findings at the molecular level. In both IO360 and DSP datasets, expression of the proliferation marker Ki-67 (MKI67) was markedly reduced upon treatment in BRCA-MUT but not in BRCA-WT tumors, at both the transcript and protein levels (**Fig. 2E, Supplementary Fig. 2E**). These findings are consistent with decreased tumor cell frequency and density by CyCIF and support preferential suppression of tumor growth in BRCA-MUT tumors as shown by clinical benefit parameters (ORR, PFS and OS) outlined above.

To determine whether the structural stability of BRCA-WT tumors reflects uniform tumor-cell persistence or emergence of distinct tumor-cell states, we examined tumor-core regions identified morphologically on H&E sections and evaluated panCK expression by CyCIF. The frequency of panCK-high tumor cells decreased in BRCA-MUT tumors and remained stable in BRCA-WT tumors after treatment (**Supplementary Fig. 2F**). However, two BRCA-WT tumors exhibited expansion of a panCK-low tumor-cell population at BX3 (**Supplementary Fig. 2G**). Comparison of matched samples revealed elevated pERK expression in panCK-low cells relative to panCK-high cells, suggesting a distinct signaling phenotype in this subset (**Supplementary Fig. 2H,I**). These findings suggest that in a subset of BRCA-WT tumors, structural stability is accompanied by the emergence of a panCK-low, pERK-high tumor-cell state rather than uniform tumor-cell persistence.

### Talazoparib modulates DNA-repair pathways and is associated with increased STING-pathway expression in BRCA-MUT tumors

We previously demonstrated that PARP inhibition induces DNA damage accompanied by activation of the cGAS–STING pathway and subsequent recruitment of CD8+ T cells in preclinical models of Brca1-deficient TNBC.^25, 27^ Therefore, we next examined whether one month of talazoparib exposure was associated with changes in expression of DNA damage response (DDR) programs and cGAS-STING-related genes. This analysis was enabled by the 55 DDR-related genes added to the IO360 panel (**Supplementary Data 1**). In BRCA-MUT but not WT tumors, genes related to DDR increased expression after talazoparib and were sustained through combination therapy (**Supplementary Fig. 3A-C**). An exception included *POLQ* expression, associated with MMEJ, which decreased following treatment of BRCA-MUT tumors (p < 0.05), potentially reflecting its association with a mitotic and/or proliferative state^37–39^, whereas no comparable change was observed in BRCA-WT (**Supplementary Fig. 3A-C**).

In parallel, genes related to a composite cGAS–STING signature increased in BRCA-MUT tumors, with relatively little to no induction in BRCA-WT tumors (**Fig. 3A-D, Supplementary Fig. 3D**). Longitudinal trajectories indicated that the transcriptional changes observed from BS to BX2 were sustained through BX3 (**Fig. 3C**, **Supplementary Fig. 3D**). Consistent with this pattern, the overlapping interferon-α response signature showed sustained induction in BRCA-MUT tumors but relative downregulation in BRCA-WT tumors across BX2 and BX3 (**Fig. 3D**). Supporting these transcriptional findings, STING protein, an upstream sensor of DNA damage-induced type I interferon signaling, showed concordant changes when measured by DSP, including decreased expression in the stroma of BRCA-WT tumors following treatment (**Fig. 3E**).

**Figure 3.**
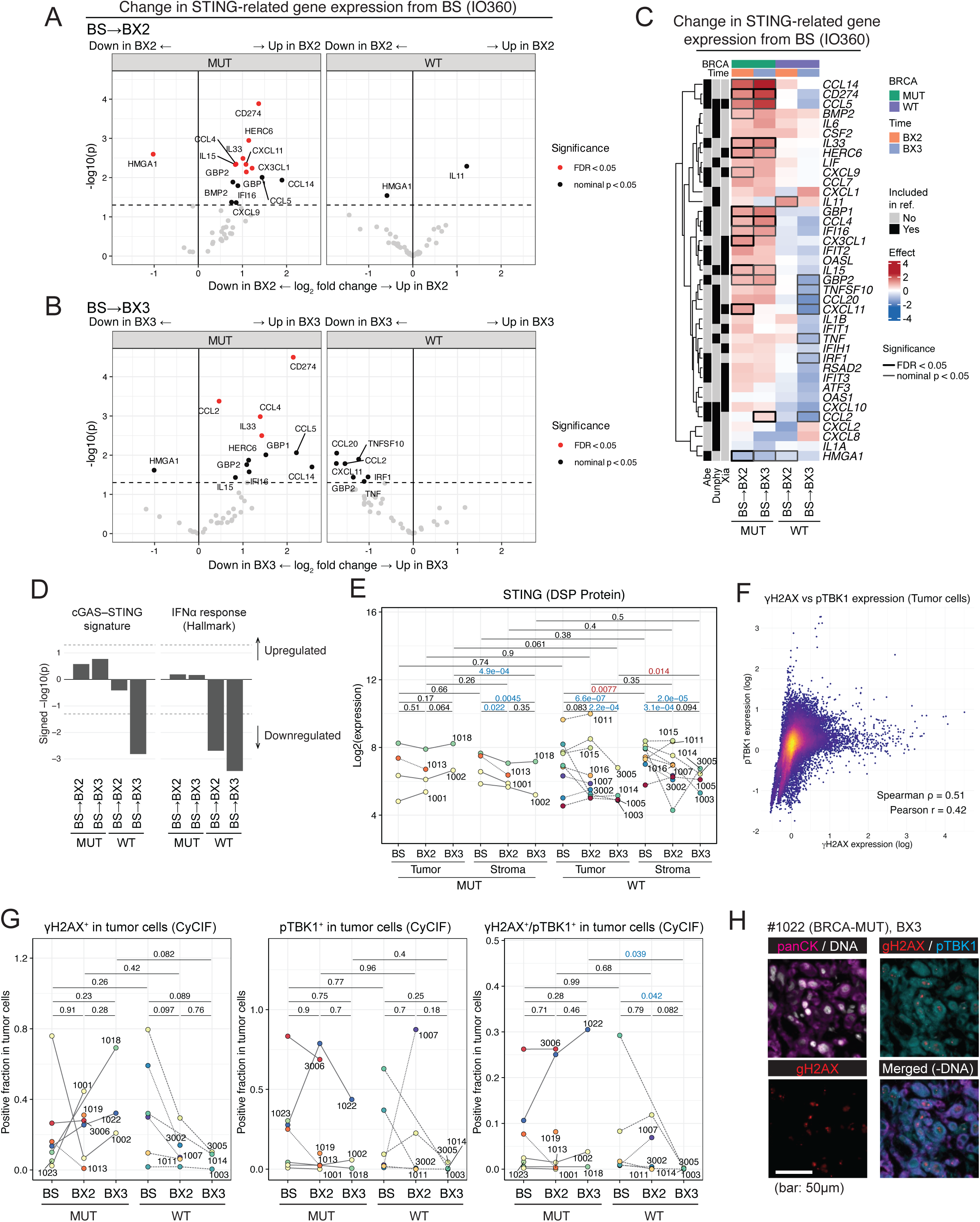
cGAS–STING signaling is attenuated in BRCA-WT tumors following PARP inhibition. (A–B) Volcano plots showing differential expression of cGAS–STING pathway genes from baseline (BS) to BX2 (A) and BX3 (B) in BRCA-MUT and BRCA-WT tumors using IO360 RNA profiling. (C) Heatmap of cGAS–STING pathway genes across patients. The signature was compiled from three published studies (annotated at left). Values represent expression changes relative to baseline. (D) Gene signature analysis of a custom cGAS–STING signature (left; same as in C) and the Hallmark IFN-α response signature (right), shown as change from baseline at BX2 and BX3 in BRCA-MUT and BRCA-WT tumors. (E) STING protein expression measured by DSP across time points, patients, and segments (tumor/stroma). (F) Expression of γH2AX and pTBK1 in tumor cells pooled across all samples measured by CyCIF. (G) Frequencies of γH2AX+, pTBK1+, and γH2AX+/pTBK1+ double-positive tumor cells measured by CyCIF. (H) Representative images of γH2AX+/pTBK1+ double-positive tumor cells (γH2AX:red; pTBK1:cyan) in CyCIF. In A and B, the y-axis shows nominal −log10(P); point color reflects a three-tier significance scheme (red: FDR < 0.05; black: nominal P < 0.05 only; grey: not significant). In C, box border denotes FDR < 0.05 (black) or nominal P < 0.05 only (grey). In D, bar height reflects signed −log10(FDR), as computed by fgsea.

We utilized CyCIF to assess γH2AX and pTBK1 as surrogate markers of DNA damage response and cGAS-STING signaling, respectively. Among tumor cells, γH2AX and pTBK1 were strongly co-expressed (**Fig. 3F, Supplementary Fig. 3E, Supplementary Fig. 4)**. The frequencies of γH2AX+ and γH2AX+/pTBK1+ tumor cells remained stable or modestly increased in BRCA-MUT tumors, whereas both populations declined in BRCA-WT tumors from BS to BX3 (p = 0.089 and p = 0.042, respectively; **Fig. 3G-H**). Consistent with these changes, mean tumor-cell γH2AX and pTBK1 expression also decreased in BRCA-WT tumors from BS to BX3 (p = 0.04 and p = 0.077, respectively; **Supplementary Fig. 3F**).

### PARP inhibition drives immune activation and suppresses proliferation in BRCA-MUT but not BRCA-WT tumors

Having established clinical evidence of PARP inhibitor-induced DNA damage and cGAS–STING activation in BRCA-MUT, but not BRCA-WT tumors, we next performed unbiased pathway-level analysis of the IO360 transcriptomic data, revealing divergent responses between the two groups of tumors (**Fig. 4A**). In BRCA-MUT tumors, proliferation-associated pathways, including E2F targets and mitotic spindle programs, were significantly downregulated relative to baseline, whereas these pathways increased in BRCA-WT tumors following treatment. In contrast, immune pathways, including interferon responses, TNFα/NF-κB signaling, and the cGAS–STING-related program, were sustained or modestly increased in BRCA-MUT tumors but were suppressed in BRCA-WT tumors, particularly at BX3.

**Figure 4.**
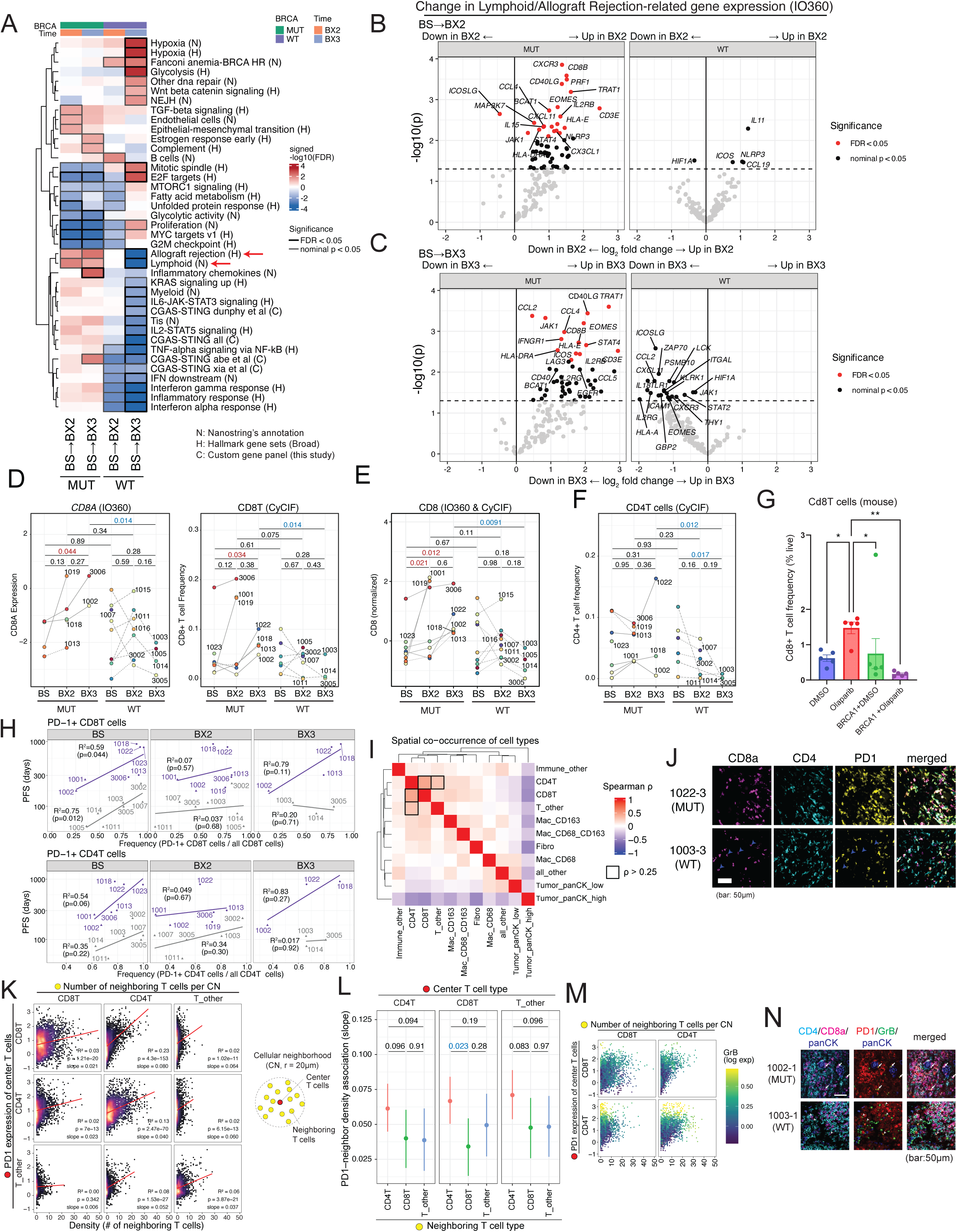
BRCA-dependent remodeling of the TME links PD-1 expression in T cells to CD4+ T cell density and PFS. (A) Gene set enrichment analysis of Hallmark (H), NanoString-annotated (N), and custom cGAS-STING (C) gene sets from baseline (BS) to BX2 and BX3 in BRCA-MUT and BRCA-WT tumors (IO360 RNA profiling). P values in A are FDR-adjusted (Benjamini–Hochberg): boxes denote significantly enriched pathways (black: FDR < 0.05; grey: nominal P < 0.05 only). (B–C) Change in Lymphoid– and Allograft Rejection-related gene expression from BS to BX2 (B) and BX3 (C) (IO360). Point color reflects a three-tier significance scheme (red: FDR < 0.05; black: nominal P < 0.05 only; grey: not significant), as in Fig. 3A,B. (D) CD8A RNA expression (IO360; left) and CD8+ T cell frequency (CyCIF; right) per patient, stratified by BRCA status and time point. (E) Integrated CD8 signal (IO360 CD8A + CyCIF CD8+ T cell frequency, z-scored and combined) per patient, stratified by BRCA status and time point. (F) CD4+ T cell frequency (CyCIF) per patient, stratified by BRCA status and time point. (G) CD8+ and CD4+ T cell frequency (percentage of live cells) in BRCA1-mutant and wild-type mouse tumors following olaparib or DMSO treatment. Bars show mean ± SEM. (H) Fraction of PD-1+ cells within CD8+ and CD4+ T cell subsets at each time point and association with progression-free survival (PFS; days, log scale). (I) Co-expression of cell state markers across CNs. Marker pairs with Spearman correlation > 0.25 are highlighted with boxes. (J) Representative images of CD4+ and CD8+ T cells and PD-1 expression in BRCA-MUT and BRCA-WT patients at BX3. (K) Association between PD-1 expression in T cells and the number of neighboring T cells within 20-µm CNs, stratified by T cell subsets across all samples. (L) Bayesian regression modeling PD-1 expression in each T cell subtype (CD8+, CD4+, T_other) as a function of local CD8+, CD4+, and T_other T cell density. Points show posterior point estimates with 95% credible intervals; brackets denote pairwise posterior comparisons between neighboring-density coefficients within each center cell type. (M) Same analysis as in K, with points colored by Granzyme B expression of center T cells (CD4+ and CD8+ T cells only). (N) Representative micrograph of GrB expression in CD4+ and CD8+ T cells from BRCA-MUT and BRCA-WT samples at BS. Arrows indicate GrB+ CD8+ T cells, while arrowheads indicate GrB+ CD4+ T cells. P values in D, E, and F are from two-sided linear mixed-effects models with patient as a random effect. P values in H are from two-sided linear regression. P values in K are from the same test, on cells pooled across all patients and samples for illustration. L shows posterior slope estimates from Bayesian mixed-effects regression.

Consistently, two independent immune pathways, including Lymphoid and Hallmark Allograft Rejection, showed sustained upregulation at both BX2 and BX3 in BRCA-MUT tumors while declining in BRCA-WT tumors. Gene-level analysis confirmed these patterns (**Fig. 4B,C**). In BRCA-MUT tumors, T cell-associated genes, including *CD8B* and *CXCR3*, encoding a receptor for the interferon-inducible chemokines CXCL9–11, were among the most strongly upregulated following treatment and remained elevated at later time points, as were the cytotoxic/effector markers *CCL4*, *IL2RB*, and *EOMES*. In contrast, these immune-related genes were downregulated in BRCA-WT tumors, consistent with the genotype-specific divergence observed at the pathway level. Patient-level analyses further supported these relationships, as shown for *CD8A* gene expression, which increased in BRCA-MUT tumors after treatment but decreased or remained largely unchanged over time in BRCA-WT tumors (**Fig. 4D**). Correspondingly, the frequency of CD8+ T cells among all cells measured by CyCIF increased in BRCA-MUT tumors after treatment but decreased or remained unchanged in BRCA-WT tumors (**Fig. 4D**). Together, these analyses demonstrate that PARP inhibition in BRCA-MUT tumors is associated upregulation of compensatory DDR gene expression programs and cGAS–STING signaling accompanied by immune activation, with suppression of proliferative programs, a pattern not observed in BRCA-WT tumors.

### DSP protein profiling confirms suppression of immune pathways in BRCA-WT tumors following PARP inhibition

Although bulk transcriptomics and immune cell frequencies relative to all cells suggest increased CD8+ T cell abundance and effector activity on treatment in BRCA-MUT tumors, these measures may be influenced by shifts in tumor cell content and reflective of the decline in total tumor cellularity over the course of treatment in BRCA-MUT tumors (**Fig 2B-D**), rather than any true change in immune cell number or activity. We therefore corroborated these pathway-and gene-level patterns using compartment-resolved (DSP) to quantify protein-level expression changes from BS to BX2 and BX3 separately within panCK-positive (tumor) and panCK-negative (stromal) regions for each BRCA subgroup (**Supplementary Fig. 5A**). Because DSP quantifies proteins independently within these compartments, it enables comparison of treatment effects with minimal influence from tumor cell purity. Across the full DSP proteome panel, protein expression tended to remain stable in BRCA-MUT tumors. In contrast, multiple immune-related proteins—including pan-immune (CD45, CD45RO), myeloid (CD14, CD68, CD163, CD11c, HLA-DR), and T cell (CD3, CD4, CD8) markers, as well as immune checkpoint proteins (PD-L1, VISTA)—were significantly downregulated over time in the stromal compartment of BRCA-WT tumors (**Supplementary Fig. 5A,B**). The reduction of these markers suggests decreased abundance of T cell and macrophage populations within the TME of BRCA-WT tumors. Concomitant downregulation of STING in BRCA-WT tumors (**Fig. 3E**) is consistent with attenuation of a key mediator of DNA damage-induced type I interferon signaling and was accompanied by reduced PD-L1 expression. Together, these protein-level changes in BRCA-WT tumors align with the transcriptional reduction of cGAS–STING and immune activation pathways observed.

### Multimodal analysis of CD8+ T cell dynamics further supports divergent immune responses in BRCA-MUT and BRCA-WT tumors

As CD8+ T cells have been shown to be a key mediator of PARPi efficacy in mouse models of Brca-deficient cancer^27^, we focused on CD8+ T cell dynamics across patients and integrated the IO360 and CyCIF datasets using LMER while controlling for patient– and platform-specific variation. This integrative analysis revealed a significant increase in CD8+ T cell frequency from baseline to BX2 and BX3 in BRCA-MUT tumors, whereas BRCA-WT tumors showed a gradual decline over time. Although this reduction was not statistically significant due to baseline variability, by BX3 all BRCA-WT tumors exhibited uniformly low CD8+ T cell levels, resulting in a significant difference between BRCA-MUT and BRCA-WT groups (**Fig. 4E**).

DSP analyses covered partially overlapping sample sets across technologies because DSP profiling requires regions with sufficient tissue area for spatial segmentation, whereas CyCIF and IO360 analyses were performed on independently available specimens. Nevertheless, protein-level changes were mostly consistent with these findings (**Supplementary Fig. 5C,D**). CD4+ T cell frequencies measured by CyCIF showed a similar pattern, with sustained levels in BRCA-MUT tumors but a significant decline in BRCA-WT tumors, again leading to a marked difference between the groups by BX3 (**Fig. 4F**). Notably, the decline in both CD8+ and CD4+ T cell abundance in BRCA-WT tumors following therapy is consistent with the absence of tumor DNA damage and cGAS–STING activation after PARPi exposure. A similar pattern was observed in our immunocompetent mouse model of Brca1-deficient TNBC, where PARPi treatment drove an increase in CD8+ T cells, whereas in BRCA1-restored TNBC there was a loss of CD8+ T cells following therapy (**Fig.4G**). Together, these findings provide clinical evidence of CD8+ T cell accumulation in BRCA-MUT tumors, whereas BRCA-WT tumors exhibit progressive loss of T cells during treatment.

### PD-1 expression in T cells is associated with clinical outcome and CD4+ T cell proximity

We next examined the relationship between T cells and PFS, prioritizing metrics normalized within specific immune compartments, including fractions of PD-1+ CD8+ or CD4+ T cells among all CD8+ or CD4+ T cells, i.e., measures not affected by changes in tumor cell content over the course of treatment in BRCA-MUT tumors, unlike immune cell frequencies expressed relative to all cells. Consistent with this distinction, the overall abundance of T cells or PD-1+ T cells as a frequency among all cells did not correlate strongly with clinical outcome (**Supplementary Fig. 6A-B**). In contrast, the fraction of PD-1+ CD8+ T cells and PD-1+ CD4+ T cells out of total CD8+ and CD4+ T cells, respectively, showed a stronger association with PFS, particularly at baseline and BX3 in BRCA-MUT tumors, whereas these relationships were weaker at BX2 and largely absent in BRCA-WT tumors (**Fig. 4H, Supplementary Fig. 6C**).

To investigate the spatial context of the PD-1+ T cells, CyCIF imaging was used to define three T cell populations: CD8+, CD4+, and CD3+CD4^−^CD8^−^ (“T_other”) cells. These T cell subsets frequently co-localized with one another, forming T cell-enriched niches distinct from non-T cell regions (**Fig. 4I,J**). Although the overall abundance of PD-1+ T cells did not change substantially during treatment (**Supplementary Fig. 6D**), we hypothesized that PD-1 expression might be influenced by the local T cell microenvironment rather than total cell abundance. We therefore examined PD-1 expression in relation to local T cell density within cellular neighborhoods (**Fig. 4K**). PD-1 expression in T cells increased with higher local T cell density, with the strongest relationship observed between PD-1 expression in CD8+ T cells and proximity to CD4+ T cells. This suggests that CD4+ T cell-rich niches may promote PD-1 expression in neighboring CD8+ T cells.

To disentangle the contributions of different T cell subsets, we modeled PD-1 expression in each T cell subtype as a function of local CD8+, CD4+, and T_other T cell density using a Bayesian hierarchical model (**Fig. 4L**; see Methods). PD-1 expression increased with the density of all three neighboring T cell subtypes across all three center cell types, with CD4+ T cell density showing the strongest association within each center subtype — significantly exceeding the association of CD8+ T cell density with PD-1 expression within CD8+ T cells (posterior comparison, p = 0.023), and directionally consistent, though not significant, in CD4+ and T_other center cells. To test whether this CD4+-dominant pattern held across clinical subgroups, we modeled group-level slopes hierarchically with partial pooling by BRCA status × timepoint group, confirming that the CD4+ T cell density coefficient was consistently larger than the CD8+ T cell density coefficient (Wilcoxon, p = 0.031; **Supplementary Fig. 6E**), a pattern also confirmed by individual-cell partial residuals (**Supplementary Fig. 6F**).

Notably, although granzyme B (GrB) expression tended to increase with higher PD-1 expression at the single-cell level in both CD8+ and CD4+ T cells, the majority of PD-1+ cells in either subset lacked detectable GrB expression (**Fig. 4M,N**). Consistent with this finding, the fraction of PD-1+/GrB+ cells was not positively associated with PFS in either CD8+ or CD4+ T cells — and in several BRCA-MUT subgroups showed a significant or near-significant negative relationship — whereas PD-1+/GrB– cells in both subsets were more prevalent in samples with longer PFS (**Supplementary Fig. 6G**). These observations suggest that tumors with favorable outcomes may be characterized by a larger pool of recently recruited or early-activated CD8+ and CD4+ T cells marked by PD-1 expression without cytotoxic differentiation, whereas tumors with fewer T cells tend to contain a higher fraction of PD-1+/GrB+ cells that may represent more terminally differentiated or exhausted states.

### Macrophage dynamics diverge between BRCA-MUT and BRCA-WT tumors

We next used CyCIF image analysis to determine immune cell changes more broadly across longitudinal samples. Two major cell-type frequency clusters emerged (**Fig. 5A, Supplementary Fig. 7A**). Cluster 1 encompassed broad immune populations, including T cells and macrophages, and their subsets such as CD8+ T cells and CD163+ macrophages (i.e., CD68–/CD163+), which increased in BRCA-MUT tumors after treatment. This increase may partly reflect reduced tumor cell abundance, which would be associated with an increased proportion of non-tumor cells, rather than *de novo* infiltration (**Fig. 2B**). Cluster 2 comprised CD4+ T cells and CD68+CD163+ double-positive macrophages. Unlike Cluster 1, Cluster 2 remained stable in BRCA-MUT tumors but declined significantly in BRCA-WT tumors (**Fig. 5A**). Because tumor purity did not change substantially in BRCA-WT tumors, this reduction likely reflects a genuine biological decrease rather than compositional bias (**Supplementary Fig. 7B,C**).

**Figure 5.**
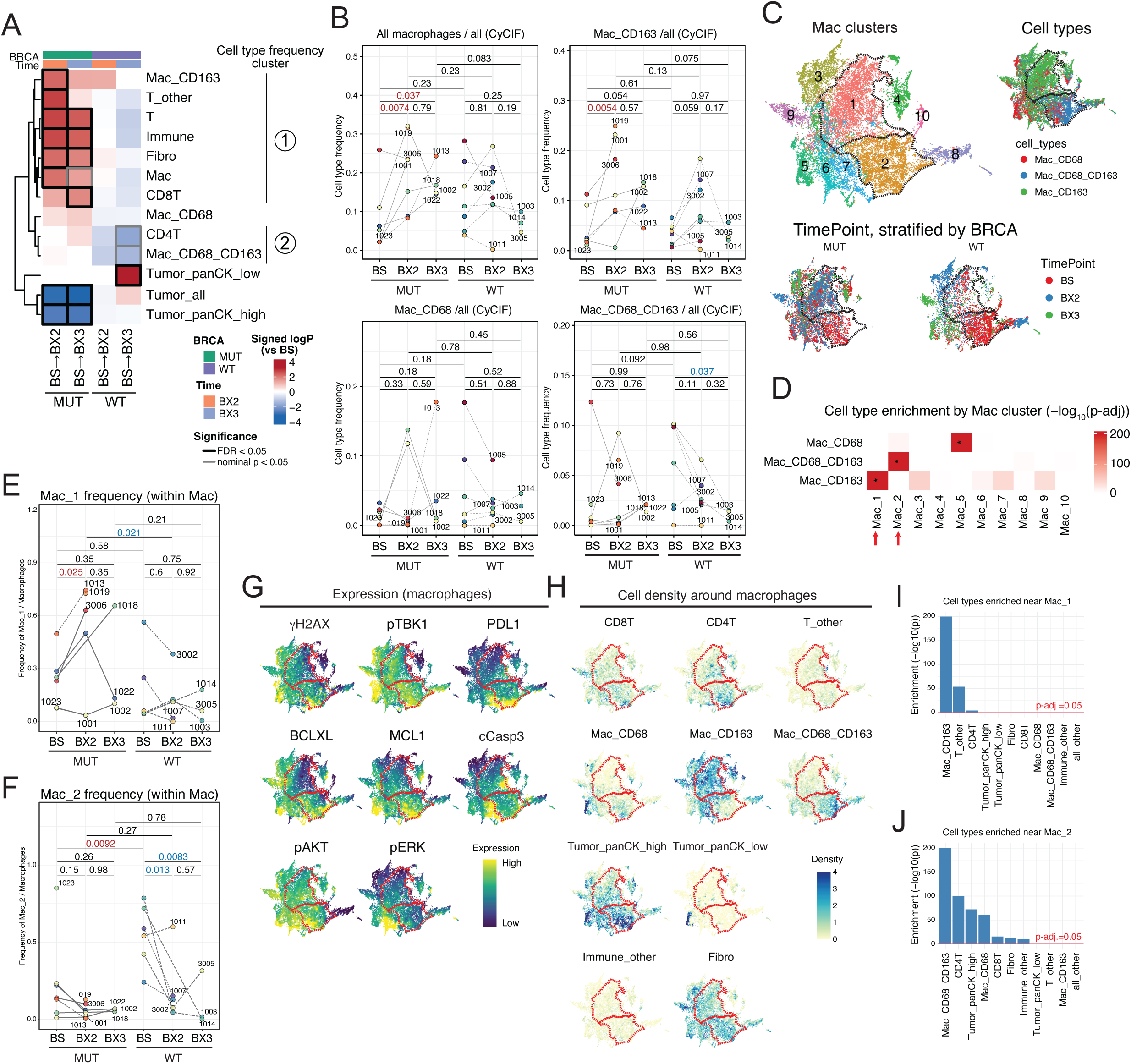
Macrophage subtypes are associated with clinical outcome and display distinct spatial distributions. (A) Heatmap of cell type frequency changes across treatment and BRCA status. Each row represents a comparison: BRCA-MUT (top), BRCA-WT (middle), and BRCA-WT vs. BRCA-MUT (bottom). Colors show signed log₁₀ p values from linear mixed-effects models (red: higher in right-hand condition; blue: higher in left). Non-tumor cell types with significant changes are clustered into two groups (Cluster 1/Cluster 2) based on temporal dynamics within each BRCA subgroup. (B) Frequency of total macrophages and three subtypes of macrophages among all cells, stratified by BRCA status and time point. (C) UMAP of pooled macrophages from all samples. Top left: macrophage clusters. Top right: color-coded by subtype (CD68+, CD163+, or CD68+/CD163+). Bottom: color-coded by time point, split by BRCA group. Dashed line highlights cluster Mac_1 and Mac_2. (D) Enrichment of macrophage subtypes in each cluster, showing overrepresentation of CD163+ and CD68+/CD163+ double-positive macrophages in Mac_1 and Mac_2, respectively. (E,F) Relative frequency of (E) Mac_1 and (F) Mac_2 macrophages among all macrophages by time point and BRCA status. (G) UMAPs of macrophages colored by expression of cell lineage (CD68, CD163) and cell state markers. (H) UMAPs of macrophages, each colored by the local count of one neighboring cell type (one UMAP per cell type). (I,J) Cell types found more frequently than expected among cells adjacent to (I) Mac_1 and (J) Mac_2 macrophages. P values are BH-adjusted for multiple comparisons in A, D, I, and J.

Building on our prior findings that PARP inhibition increases macrophage infiltration in Brca1-deficient mouse tumors^25^, we examined CD68+ and/or CD163+ macrophages in more detail. Overall macrophage frequency rose over time (**Fig. 5B**), consistent with Cluster 1, but individual subsets diverged: CD163+ macrophages increased sharply, primarily between BS and BX2 in BRCA-MUT tumors, and were variable in BRCA-WT tumors at BX2 and BX3, whereas CD68+CD163+ macrophages were generally stable in BRCA-MUT tumors, but decreased in most BRCA-WT tumors (**Fig. 5B**). By contrast, CD68+ macrophages showed no significant changes.

### PARP inhibition remodels tumor–stroma architecture and macrophage distribution

We then trained a simple classifier based on the composition of neighboring cells within a 20-µm radius, which accurately separated tumor and stromal regions and aligned well with, though refined beyond, the approximate boundaries defined by local cell composition (**Supplementary Fig. 8A**). This analysis complements that shown in **Fig. 2C**, where tumor cell density highlighted structural disruption in BRCA-MUT but not BRCA-WT tumors. The tumor–stroma classification adds another dimension by explicitly distinguishing these compartments at the single-cell level and capturing how they intermix. In particular, the Delaunay mixing score, which quantifies this intermingling, increased over time in BRCA-MUT tumors—indicating progressive disintegration of tumor–stroma boundaries—whereas it remained low and stable in BRCA-WT tumors (**Supplementary Fig. 8B**). This tumor–stroma classification framework provided a spatial reference for subsequent analyses, enabling cell-type–specific localization, quantification of boundary dynamics, and interpretation of immune and stromal remodeling relative to tumor and stromal compartments.

We then used this method to reveal that macrophage subsets exhibited distinct spatial distributions. CD68+ macrophages were enriched within tumor-core regions (defined computationally as areas containing ≥10 physically interacting tumor cells), whereas CD163+ macrophages were predominantly stromal. CD68+CD163+ macrophages exhibited an intermediate distribution, bridging tumor and stromal regions—consistent with findings from an independent study (**Supplementary Fig. 8C,D**).^40^ Several other immune and stromal cell types also showed clear tumor-stromal spatial distribution (**Supplementary Fig. 8E**).

### Macrophage –T cell interactions shape local immune niches

To explore macrophage heterogeneity, we pooled macrophages across samples and projected them into a UMAP based on lineage and cell state marker expression (**Fig. 5C**, **Supplementary Fig. 8F**). This analysis revealed distinct distributions of CD68+, CD163+, and CD68+CD163+ subsets across BRCA status and treatment time points. Leiden clustering identified ten macrophage clusters (**Fig. 5C top left**), among which Mac_1 and Mac_2 were enriched for CD163+ and double-positive macrophages, respectively (**Fig. 5D**).

Mac_1, the largest cluster, increased in frequency after PARPi treatment in BRCA-MUT tumors, consistent with the rise of CD163+ macrophages (**Fig. 5E**). In contrast, Mac_2 was most abundant at baseline and showed a progressive decline in both BRCA-MUT and BRCA-WT tumors; this trend was particularly pronounced in BRCA-WT tumors, consistent with the marked loss of double-positive macrophages (**Fig. 5F**). Mac_2 occurred more frequently in tumor regions (**Supplementary Fig. 8G**). This pattern supports distinct spatial niches for macrophage subsets within the TME.

Phenotypically, Mac_1 macrophages showed low expression across all cell state markers assessed, whereas Mac_2 macrophages exhibited higher PD-L1 levels (**Fig. 5G**), suggesting a more activated phenotype. We then evaluated their relationship with T cells. Neighborhood analysis further revealed that Mac_1 macrophages were most frequently associated with T_other cells and, to a lesser extent, CD4+ T cells, whereas Mac_2 macrophages showed strong co-localization with CD4+ T cells (**Fig. 5I,J**). Notably, CD68+CD163+ macrophages and CD4+ T cells belonged to the same cell type frequency cluster (Cluster 2), which was most abundant at baseline and declined in BRCA-WT tumors (**Fig. 5A**). Although these associations do not establish a causal link to the longitudinal dynamics, their spatial co-localization suggests that these cell types form shared immune niches within the TME. Other macrophage clusters were not enriched for CD68+CD163+ macrophages and did not exhibit similar temporal dynamics to Mac_2 (**Fig. 5D**, **Supplementary Fig. 8H**).

This macrophage-centric analysis provided further evidence that immune activity is sustained or stable in BRCA-MUT tumors but declines in BRCA-WT tumors during PARP inhibition. In BRCA-MUT tumors, CD163+/PD-L1-low macrophages enriched in Mac_1 that interact with T_other and CD4+ T cell subsets increased. In contrast, BRCA-WT tumors lost a peri-tumoral CD68+CD163+ macrophage subset that co-localized with CD4+ T cells in regions of higher PD-L1 expression. Across both BRCA groups, Mac_2, enriched in PD-L1 expression, declined over time (**Fig. 5F**), suggesting limited opportunity for effective PD-L1 blockade despite PARPi-induced immune remodeling.

### Distinct CD163+ macrophage-rich CNs that expand after PARP inhibition show different T cell associations

Effective anti-tumor immunity depends not only on immune cell abundance but also their spatial organization with tumor cells. To quantitatively capture these contexts, we applied cellular neighborhood (CN)–level analysis, in which each neighborhood—defined as cells within a 20-µm radius—is treated as an analytical unit. Each CN is characterized by its local cell-type composition together with aggregated cell-state marker expression, enabling direct comparison of neighborhood states within and across samples. We used this approach to identify therapy-responsive niches and determine which neighborhoods persist or collapse across BRCA genotypes after treatment — these patterns that cannot be resolved by analyzing cell types in isolation.

UMAP projection followed by Leiden clustering identified 17 distinct TME clusters (**Fig. 6A**, **Supplementary Fig. 9A,B**). Cell-type frequencies and marker expression varied widely among clusters, reflecting heterogeneous tumor, immune and stromal compositions (**Fig. 6B,C, Supplementary Fig. 9C,D**). TME_1 comprised CNs in which neighboring cells were >80% tumor cells, whereas the remaining clusters contained fewer tumor-dominant CNs (**Supplementary Fig. 9E**). TME_1 decreased significantly in BRCA-MUT tumors following treatment (**Supplementary Fig. 9F**), consistent with our prior analysis of tumor architecture (**Fig. 2B-E**).

**Figure 6.**
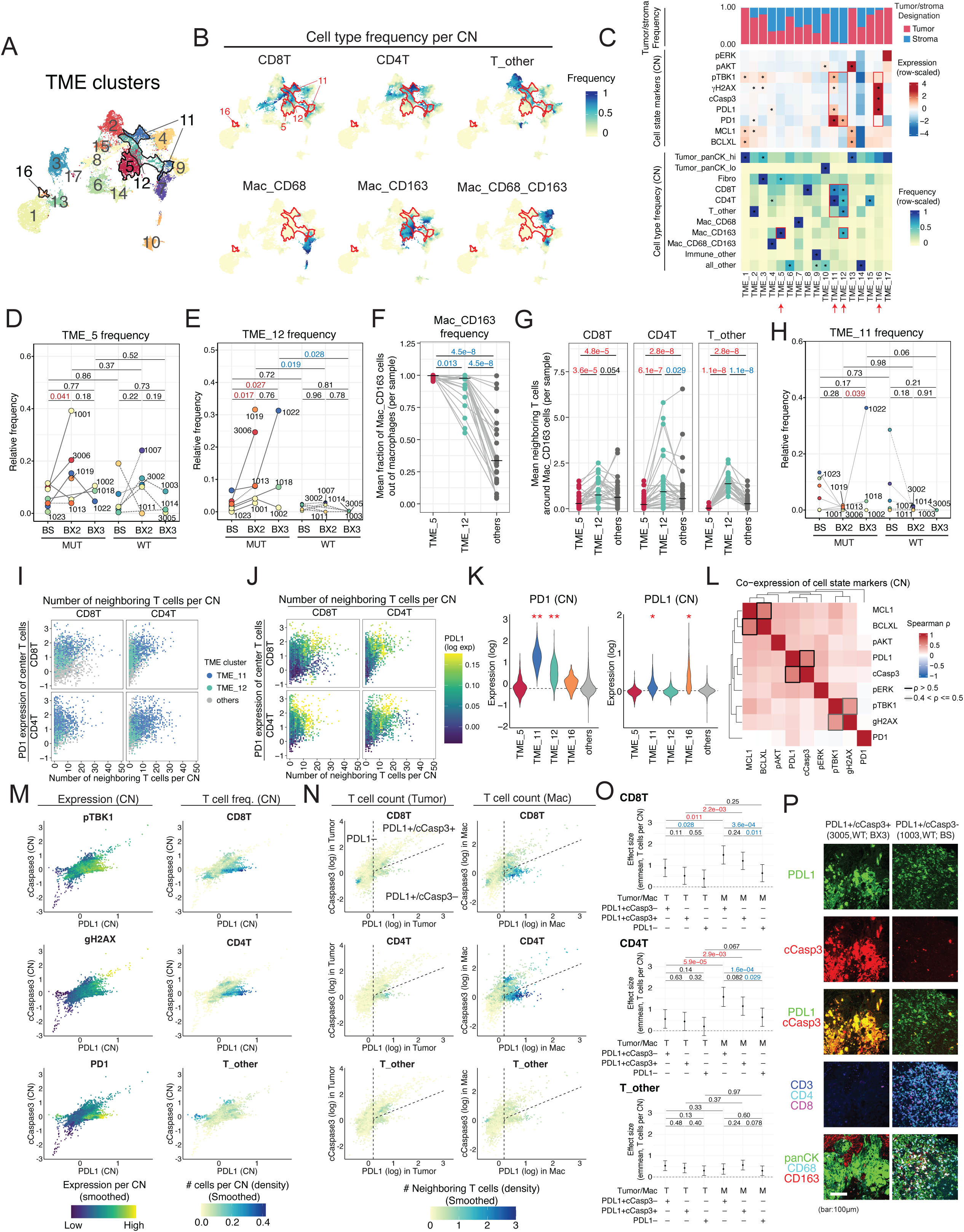
Clustering of CNs reveals immunologically distinct PD-L1+ microenvironments. (A) UMAP of cellular neighborhoods (CNs) clustered into 17 groups based on cell type frequencies and state marker expression. Boundaries of TME_5, TME_11, TME_12, and TME_16 are highlighted. (B) UMAPs colored by the frequency of T cell and macrophage subsets within CNs. (C) Heatmaps summarizing cluster features: tumor–stroma designation frequency (top), overexpressed proteins (middle), and overrepresented cell types (middle). Asterisks denote one-sided Wilcoxon rank-sum test (in-cluster vs. out-of-cluster values), BH-adjusted. (D,E) Relative frequency of TME_5 (D) and TME_12 (E) among all CNs, stratified by BRCA status and time point. Lines connect longitudinal samples from the same patient. (F) Mean CD163+ macrophage density in TME_5 and TME_12 versus other CNs. (G) Mean T cell density surrounding CD163+ macrophages in TME_5 and TME_12 versus other CNs. (H) Frequency of TME_11 among all CNs, stratified by BRCA status and time point. (I,J) Same 2×2 subset (CD4+/CD8+) of the density-vs-PD1 relationship shown in Fig. 4K, with points colored by TME_11/TME_12 membership (I) or, as a representative example, CN-level PD-L1 expression (J). (K) Expression of selected proteins in CNs belonging to four TME clusters (TME_5, TME_11, TME_12, TME_16) and all remaining CNs combined. (L) Pairwise Spearman correlation of cell state marker protein expression at CN-level. (M) CN-level expression of PD-L1 and cleaved caspase-3 (cCasp3), colored by protein expression (left) or by the number of neighboring T cells (right). (N) Single-cell expression of PD-L1 and cCasp3 in tumor cells (left) and macrophages (right). Each point represents an individual cell and is colored by the number of neighboring T cells (CD8+, CD4+, or T_other). Dashed lines divide plots into PD-L1+/cCasp3–, PD-L1+/cCasp3+, and PD-L1– regions. (O) Linear mixed-effects model coefficients summarizing differences in neighboring T cell density across the PD-L1/cCasp3-defined regions shown in (N). (P) Representative micrographs showing PD-L1+/cCasp3+ (left) and PD-L1+/cCasp3– (right) macrophage–T cell microenvironments. P values are BH-adjusted for multiple comparisons in C, F, G, and K; P values in I, J, and N are from cell-level regression, shown for illustration only (see Methods and Fig. 4L for the patient-level treatment); pairwise contrasts in O are nominal P values, uncorrected.

Among the 17 TME clusters, TME_5 and TME_12 increased after PARPi treatment specifically in BRCA-MUT tumors (**Fig. 6D,E**). Both clusters were characterized by a high abundance of CD163+ macrophages. In TME_5, macrophages were strongly enriched for the Mac_CD163 subtype, accounting for nearly all macrophages within the neighborhood—substantially higher than the ∼55% average across all CNs (**Fig. 6F**). Despite sharing CD163+ dominance, the two clusters differed sharply in local T cell content: CD163+ macrophages in TME_5 had significantly fewer neighboring CD8+, CD4+, and T_other T cells than the average CN (all P<0.0001), whereas those in TME_12 were preferentially surrounded by T cells, exceeding TME_5 for all three subtypes and exceeding the average CN as well for CD4+ and T_other T cells (**Fig. 6G**). The temporal dynamics of the two clusters also differed. TME_5 increased transiently at BX2 and declined by BX3 (**Fig. 6D**), closely mirroring the trajectory of Mac_1 macrophages, which were enriched for CD163+ macrophages (**Fig. 5D,E**). In contrast, TME_12 increased from baseline in both BX2 and BX3 in a small subset of BRCA-MUT tumors but were largely absent in BRCA-WT tumors (**Fig. 6E**). These findings indicate that CD163+ macrophages participate in distinct spatial immune niches with different T cell associations and temporal responses to therapy.

### TME_11 vs TME_12 represent distinct T cell niches with divergent temporal dynamics

To further characterize T cell-enriched spatial niches, we next examined clusters in which T cells constituted the majority of cells. TME_11 and TME_12 were the only two clusters in which T cells comprised more than 50% of all cells, including CD4+, CD8+, and T_other subsets (**Supplementary Fig. 9G**). Although both represented T cell–aggregate niches, their temporal dynamics differed markedly. TME_11 was abundant at baseline but declined sharply after treatment in most tumors, whereas TME_12 was still detectable and appeared to increase in a small number of evaluable BRCA-MUT tumors (**Fig. 6H**).

TME_11 contained denser T cell aggregates, higher PD-1 expression, and a greater proportion of CD4+ T cells compared with TME_12 (**Fig. 6I**). CNs within TME_11 showing elevated PD-L1 expression mapped precisely to the densest CD4+/CD8+ aggregates (**Fig. 6J,K**). γH2AX and pTBK1, were also higher in these densely clustered T cell regions within TME_11 (**Supplementary Fig. 10A,B**). PD-L1, γH2AX, and pTBK1 expression at cell type levels confirmed similar trends; TME_11 expressed these markers higher than in TME_12 (**Supplementary Fig. 10C-F**). Notably, across TME clusters, PD-L1 expression was highest in macrophages, particularly CD68+CD163+ double-positive macrophages, consistent with Mac_2 (**Supplementary Fig. 10C, Fig. 5G,H**). In fact, TME_12 showed among the lowest pTBK1 levels, highlighting fundamentally distinct immune contexts (**Supplementary Fig. 10B,E**).

Importantly, PD-L1 expression was substantially higher in TME_11 and was largely restricted to macrophages. In contrast, TME_12 lacked appreciable PD-L1 expression. Thus, although CD163+ macrophages in TME_12 were embedded within dense T cell aggregates, their association with T cells occurred in the absence of PD-L1 expression. These observations suggest that TME_12 represents a distinct immune niche in which macrophage–T cell interactions are mediated by PD-L1-independent mechanisms.

### PD-L1+ niches separate into immune-engaged and immune-excluded states

Within T cell-rich neighborhoods, the highest PD-1 expression and densest T cell aggregates were associated with elevated PD-L1, γH2AX and pTBK1 levels corresponding largely to TME_11, a niche most abundant at baseline that declined sharply following treatment (**Fig. 6I,J, Supplementary Fig. 10A**). However, when all CNs were considered, γH2AX and pTBK1 were strongly correlated with each other but showed little correlation with PD-L1. Instead, PD-L1 expression correlated most strongly with cleaved caspase-3 (cCasp3), corresponding largely to TME_16 (**Fig. 6L, Supplementary Fig. 10G**), suggesting that PD-L1 expression across the tissue may arise in at least two distinct cellular contexts: one cCasp3-high (TME_16) and one cCasp3-low (TME_11).

When γH2AX and pTBK1 expression were compared at the CN level, the two markers showed a strong positive correlation, with CD4+ and CD8+ T cells enriched in γH2AX+/pTBK1+ neighborhoods (**Supplementary Fig. 10H**). At the single-cell level, pTBK1+/γH2AX+ macrophages were surrounded by the highest densities of CD4+ and CD8+ T cells, whereas this association was not observed around tumor cells (**Supplementary Fig. 10I,J**).

At the CN level, PD-L1 and cCasp3 were strongly correlated overall. However, a subset of neighborhoods showed high PD-L1 expression despite low cCasp3 levels. These PD-L1+/cCasp3– CNs exhibited elevated pTBK1 and PD-1 and were enriched for CD4+ and CD8+ T cells, largely corresponding to TME_11 (**Fig. 6M**). Importantly, PD-L1+/cCasp3+ neighborhoods were largely devoid of CD4+ and CD8+ T cells, corresponding largely to TME_16, where tumor cells comprised the majority of cells and macrophages also expressed PD-L1, indicating that PD-L1 expression frequently occurs in apoptotic, T cell–poor regions rather than within active T cell aggregates.

These patterns were confirmed at the single-cell level: PD-L1+/cCasp3– tumor cells and macrophages showed the strongest T cell enrichment, with CD4+ T cells particularly enriched near PD-L1+ macrophages, whereas PD-L1+/cCasp3+ cells exhibited minimal T cell contact (**Fig. 6N-P**).

### Overall conceptual model

Consistent with preclinical studies, BRCA-MUT tumors showed evidence of DNA damage-mediated cGAS–STING activation following PARP inhibition, accompanied by the emergence of a T cell-rich niche (TME_12) containing both CD4+ and CD8+ T cells, that was also detected at later treatment timepoints. This niche was characterized by PD-1+/PD-L1– T cells together with CD163+ macrophages that were also detected at each evaluable treatment timepoint. While the presence of CD163+ macrophages may restrain full T cell activation, their largely PD-L1–negative state provides limited opportunity for PD-L1 blockade to augment immune activity.

In contrast, BRCA-WT tumors did not show evidence of PARP inhibitor–induced cGAS–STING activation. PD-L1+ T cell-containing niches present at baseline were largely lost after treatment in both BRCA-MUT and BRCA-WT tumors. In BRCA-WT tumors, this occurred in parallel with reduced STING signaling and declining CD8+ T cell abundance. These changes reflect progressive immune attenuation in this homologous recombination repair-competent, PARPi-insensitive setting. Residual PD-L1+ regions in BRCA-WT tumors were predominantly associated with apoptotic, T cell-poor neighborhoods rather than immune-engaged niches.

These findings show that the spatial organization of T cells and their cooperation with macrophages—rather than PD-L1 expression alone—define immune-engaged niches that remain after PARP inhibition. Such niches arise preferentially in BRCA-MUT tumors where PARP inhibition generates immunogenic DNA damage and sustained immune engagement. In contrast, PD-L1+ regions in BRCA-WT tumors largely reflect apoptotic, immune-excluded microenvironments and therefore represent unlikely targets for effective PD-1/PD-L1 checkpoint blockade. Collectively, the data suggest that immune checkpoint blockade is unlikely to augment the effects of PARPi monotherapy in BRCA-MUT tumors, and that PARP inhibition is unlikely to further sensitize to immune checkpoint blockade in BRCA-WT tumors. A conceptual summary is shown in **Fig. 7**.

**Figure 7.**
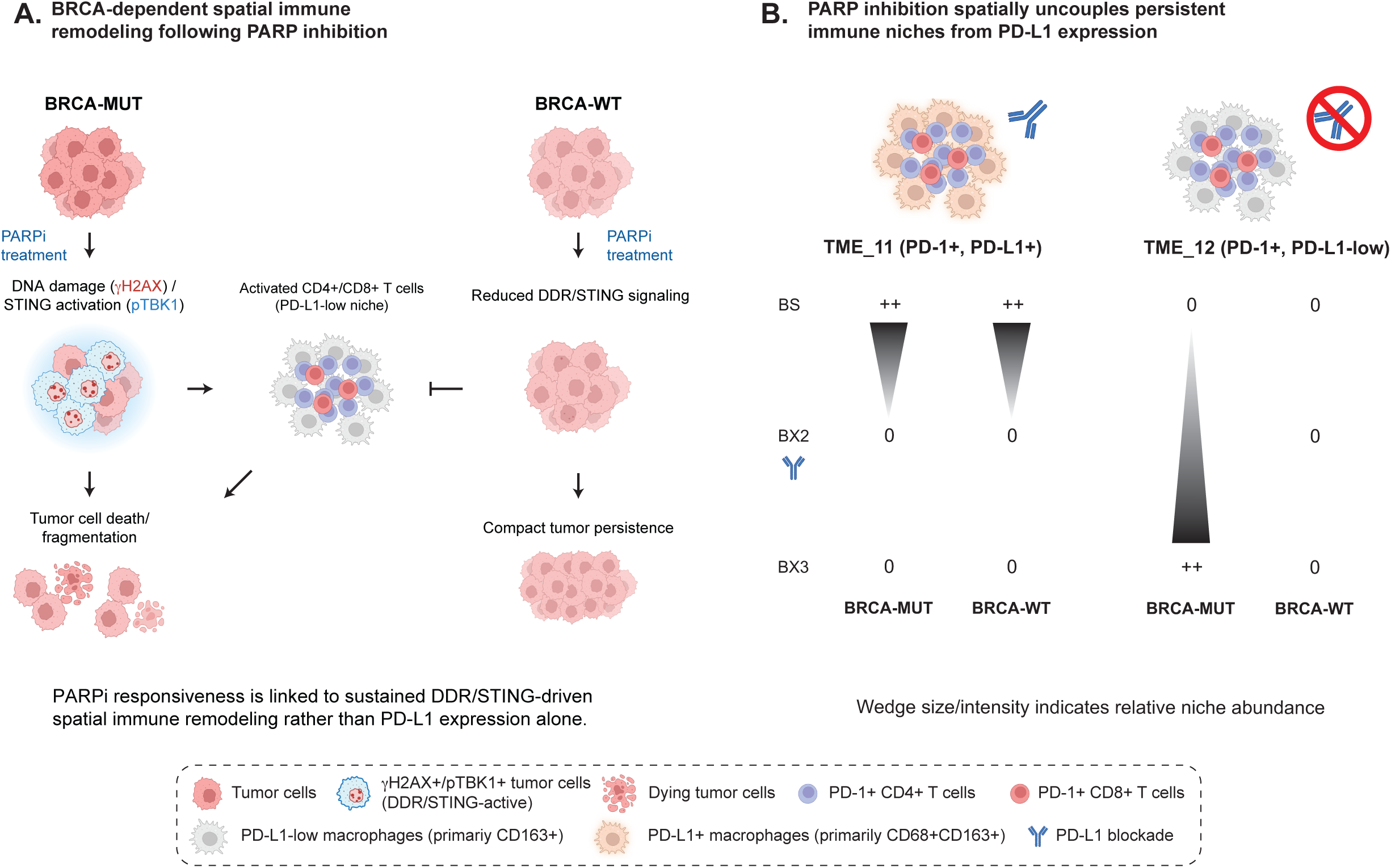
Overall conceptual model. (A) Conceptual model of BRCA-dependent spatial immune remodeling following PARP inhibition. In BRCA-MUT tumors, PARP inhibition induces DNA damage and sustained DDR/STING signaling (represented by γH2AX+/pTBK1+ tumor cells), which is associated with the emergence of activated (PD-1+) CD4+/CD8+ T cell–rich, PD-L1-low immune niches and tumor cell depletion. In contrast, BRCA-WT tumors exhibit reduced DDR/STING signaling following treatment, fail to establish durable immune-engaged niches, and maintain compact tumor architecture. These findings suggest that PARPi responsiveness is linked to sustained DDR/STING-driven spatial immune remodeling rather than PD-L1 expression alone. (B) Conceptual model of treatment-associated dynamics of distinct T cell–rich niches. TME_11 represents a transient PD-1+, PD-L1+ niche present at baseline that is depleted during treatment in both BRCA-MUT and BRCA-WT tumors. In contrast, TME_12 represents a persistent PD-1+, PD-L1-low niche that emerges selectively in BRCA-MUT tumors following treatment (based on a limited number of evaluable samples at later timepoints). These findings suggest that PARP inhibition spatially uncouples persistent immune-engaged niches from PD-L1 expression, thereby limiting opportunities for effective PD-L1 blockade despite ongoing immune remodeling. Wedge size and shading indicate the relative abundance of each niche across treatment timepoints.

## Discussion

PARP inhibitors are clinically effective in BRCA-MUT breast cancer, and preclinical studies have suggested that PARPi-induced activation of cGAS–STING signaling may enhance antitumor immunity, providing the initial rationale for TALAVE, as well as other studies in which PARP inhibition has been combined with immune checkpoint blockade. The combination of talazoparib plus avelumab was generally well tolerated, with a safety profile dominated by expected hematologic and gastrointestinal toxicities consistent with known PARPi class effects, and demonstrated substantial activity in *BRCA1/2*-mutant breast cancer.

In this work, we integrated genomic, transcriptomic, and spatial proteomic profiling of serial tumor biopsies collected across treatment time points to define how PARP inhibition, alone or in combination with PD-L1 blockade, remodels the tumor immune microenvironment. Because BX3 biopsies were limited in number and the study lacked a contemporaneous arm including patients who continued only on PARPi monotherapy, changes observed at BX3 cannot be definitively attributed to the addition of PD-L1 blockade as opposed to continued PARPi exposure alone, so that combination-phase findings must be interpreted with this caveat.

In BRCA-MUT tumors, PARP inhibition induced heterogeneous, but measurable immune activation accompanied by structural remodeling, including marked loss and architectural disruption of tumor cells. STING and interferon signaling were upregulated, and γH2AX/pTBK1 co-expression localized to T cell-rich regions. Our results are consistent with prior observations in murine models, where PARP inhibition induces innate immune activation selectively in Brca1-deficient tumors. This is important because the induction of cGAS–STING and interferon-α response signatures shown here exclusively in BRCA-mutated breast tumors provides clinical evidence that PARP inhibition generates immunogenic DNA damage in HRD contexts. This supports a mechanistic model in which unrepaired DNA fragments activate cytosolic DNA sensing pathways, driving downstream immune engagement.

Spatial analyses further revealed that PARP inhibition reshaped the organization of immune niches. A T cell-rich neighborhood characterized by PD-1+ T cells and CD163+/PD-L1-low macrophages (TME_12) emerged and remained detectable following treatment, whereas a distinct PD-1+/PD-L1+/cCasp3– T cell-dense niche (TME-11) was largely depleted. This indicates a therapy-induced spatial uncoupling of PD-1 and PD-L1; PD-1 expression increased in T cells, while PD-L1+ niches that might support PD-1/PD-L1 interactions were lost. Residual PD-L1 expression was frequently confined to apoptotic, cCasp3+ tumor regions lacking T cell infiltration, consistent with non-productive, stress-associated upregulation.

A key organizing principle emerged from single-cell spatial modeling: PD-1 expression in CD8+ T cells was most strongly associated with the density of neighboring CD4+ T cells rather than with PD-L1 expression in surrounding cells. Among the three T cell subsets, this CD4+ T cell density association significantly exceeded the association with a cell’s own subtype density specifically in CD8+ T cells, with the same direction observed, though not reaching significance, in CD4+ and T_other T cells. Consistent with this finding, patients whose tumors contained CD4-rich microenvironments exhibited longer progression-free survival, suggesting that CD4+ T cell-supported niches sustain CD8+ cytotoxic T cell engagement following PARP inhibition. These findings are consistent with prior studies demonstrating that CD4+ T cells, including T follicular helper subsets, coordinate antitumor immune responses in part through interactions with B cells and other immune populations.^41^

Complementary spatial mapping of cellular neighborhoods revealed that CD163+ macrophage–T cell interactions in a specific spatial niche (TME_12) — but not the separate, T cell–poor CD163+-dominant niche (TME_5) — became the predominant immune-supporting compartment after treatment in BRCA-MUT tumors, but not in BRCA-WT disease. In these regions, CD163+ macrophages, although largely PD-L1– and expressing low pTBK1, clustered tightly with PD-1+ T cells, forming a PD-L1-independent, viable immune niche that remained detectable even as PD-L1+ neighborhoods were lost. In contrast, PD-L1+/cCasp3+ apoptotic regions were consistently depleted of T cells, highlighting functional separation between immune-engaged versus immune-excluded PD-L1 states.

In summary, in BRCA-MUT tumors, although immune activation is induced, it is reorganized into PD-L1–independent niches in which T cell activity is most strongly associated with CD4+ T cell proximity and CD163+ macrophage interactions rather than with PD-L1 expression. Therefore, spatial immune niches and not PD-L1 expression alone, define productive immune engagement after PARP inhibition. The persistence of CD4+/CD8+/macrophage cooperative niches in BRCA-mutated tumors show the importance of spatially organized immune microenvironments. These niches appear to be sustained by PARP-induced immunogenic DNA damage and may serve as functional biomarkers of response that is beyond PD-L1 expression.

The PD-L1-low state of CD163+ macrophages in TME_12 indicates that the addition of PD-1 or PD-L1 blockade is unlikely to augment anti-tumor immunity in BRCA-MUT tumors and that patients with PARPi-responsive tumors are unlikely to benefit substantially from the addition of immune checkpoint blockade. Indeed, the median PFS we observed in the TALAVE BRCA-MUT cohort (7.51 months) is similar to that observed with talazoparib alone in the EMBRACA study (8.6 months) in a comparable population.^17^ Although the ORR of 80%, the percentage of patients on treatment > 2 years (25%) and median OS (50.72 months) in TALAVE was higher than expected based on results reported in EMBRACA (ORR 62.6%; 13.2% on treatment > 2 years; median OS 19.3 months),^17, 19^ the small sample size and absence of randomization does not allow attribution of these endpoints to the addition of avelumab. Finally, in a larger randomized phase II study of olaparib with or without atezolizumab in BRCA-mutated advanced breast cancer, the addition of atezolizumab did not improve PFS or OS, supporting limited additional clinical benefit from PD-L1 blockade in this setting.^33^

Notably, although TME_11, enriched in PD-L1-high cells, was most abundant at baseline and generally declined after treatment, whereas TME_12, enriched in PD-L1-low cells, increased at BX2 and BX3, two patients who responded with exceptional durability (patients 1018 and 1022) with available post-treatment tissue retained relatively high frequencies of both TME_11 and TME_12 at BX3; post-treatment samples were not available for the third exceptional responder (patient 1023). This persistence of immune-engaged niches in patients with extremely prolonged clinical benefit supports an association between sustained spatially organized T cell–macrophage interactions and favorable outcome. It is possible that patients in whom TME_11 persisted at BX3 derived greater benefit from the addition of avelumab, although the small sample size precludes firm conclusions. Notably, the fraction of PD-1+ cells within both CD4+ and CD8+ T cell compartments was consistently associated with longer PFS in BRCA-MUT patients at both baseline and BX3 (**Fig. 4H**), suggesting that preservation of a PD-1+ T cell state may help identify tumors capable of sustaining immune-engaged niches and, potentially, exceptional clinical benefit. Larger studies will be required to determine whether this feature distinguishes TME_11 persisters from non-persisters and predicts benefit from PD-1/PD-L1 blockade.

Nonetheless, for the majority of BRCA-MUT tumors, where TME_11 did not persist, enhancement of the immunologic effects of PARP inhibition may instead require combination strategies that enhance CD4+ T cell activity and reprogram PD-L1– macrophages, i.e., the CD163+ macrophages (Mac_1) that expanded selectively in BRCA-MUT tumors after treatment and participated in the dominant post-treatment immune niche. Because the CSF1–CSF1R axis promotes macrophage polarization toward an immunosuppressive state^42–45^, targeting this pathway may enhance T cell infiltration and activation. Building on these findings and prior preclinical work^25^, we are pursuing a clinical strategy combining PARP inhibition with macrophage-directed therapy using axatilimab, an anti-CSF1R antibody, to alleviate macrophage-mediated T cell suppression (NCT06488378). Future work should also determine whether these macrophages express alternative immune checkpoints that further constrain T cell activity.

In contrast, BRCA-WT tumors—despite receiving the same treatment—exhibited minimal DNA damage, loss of STING and interferon signaling, depletion of CD8+ T cells, and collapse of PD-L1+ T-cell–containing niches into apoptotic, T cell-poor regions. These features are consistent with homologous recombination repair competence and PARPi insensitivity and provide a mechanistic explanation for the lack of meaningful synergy between PARP inhibition and PD-1/PD-L1 blockade in BRCA-WT disease. Taken together, these findings suggest that PD-L1 blockade may offer limited additional benefit when added to PARP inhibition through distinct mechanisms in each genetic context.

Although immune remodeling differs fundamentally by BRCA status after PARP inhibition, i.e., BRCA_MUT tumors showed sustained CD8+ T-cell accumulation, whereas BRCA_WT tumors exhibited progressive T-cell loss, it is possible that BRCA_WT tumors may have demonstrated productive immune modulation had patients in Cohort 2 been treated with effective therapy. Therefore, comprehensive assessment of spatial immune niches and their PD-L1-dependence after chemotherapy or antibody drug conjugate exposure is warranted in BRCA-WT TNBC,^46^ which may help optimize use of immunotherapy combinations.

Across datasets, the increase in CD8+ T cells observed in BRCA-MUT tumors was modest and variable. This likely reflects a balance between DNA damage-induced immune activation, potential cytotoxic effects of PARP inhibition on T cells themselves, including PARP1 trapping-induced DNA damage and apoptosis, recently shown to directly impair T cell fitness during PARPi treatment^47^, or intrinsic vulnerability of T cells to sustained STING signaling, which can trigger interferon-associated ER stress and apoptotic pathways within T cells in addition to their activation^48–50^. Additionally, shifts in immune cell proportions following tumor regression, may have resulted in only partial and heterogeneous expansion of CD8+ T cells during treatment.

Additionally, the magnitude of T cell responses was more limited than those observed in Brca1-deficient TNBC murine models possibly reflecting species differences, sampling timepoints, or, in hormone receptor-positive tumors, immune exclusion associated with estrogen receptor signaling.^40^ These findings highlight opportunities to combine PARPi with strategies tailored to the underlying tumor immune context, including approaches that counter estrogen receptor-associated immunosuppression in hormone receptor-positive disease, modulate macrophage polarization, or sustain interferon activity without inducing T cell loss. Thus, a central clinical challenge is how to enhance and stabilize the immune-engaged niches that arise in BRCA-MUT tumors under PARP inhibition. Our data suggest that these niches are defined not by PD-L1 expression, but by CD4+ T cell–supported and macrophage-associated spatial organization, providing a framework for developing combination therapies that more effectively sustain antitumor immunity.

## Materials and Methods

### Study Design and Patient Population

TALAVE (NCT03964532) was a multi-institutional, open-label phase I/II study that enrolled two cohorts of patients with histologically confirmed advanced breast cancer not amenable to curative treatment with surgery or radiation therapy. Cohort 1 included patients with HER2-negative breast cancer harboring somatic or germline *BRCA1/2* mutations, whereas cohort 2 comprised patients with TNBC without known somatic or germline *BRCA1/2* mutation. All patients had measurable disease per RECIST 1.1 criteria.^48, 49^ Patients previously treated with a PARPi, or those who experienced disease progression during the first six months of prior anti-PD-1 or anti-PD-L1 therapy, were excluded.

All participants had biopsy-accessible disease. Participants were required to undergo research biopsies at baseline (within four weeks before Cycle 1, Day 1), following talazoparib induction monotherapy (prior to Cycle 2, Day 1), and during combination therapy (prior to Cycle 3, Day 1). Patients who derived clinical benefit, defined as RECIST 1.1-confirmed partial or complete response at any point during treatment, or stable disease for at least 24 weeks, followed by disease progression, underwent a fourth and final research biopsy at the time of disease progression. This study complied with all relevant ethical regulations, including the Declaration of Helsinki, and was approved by the MedStar Georgetown University Hospital Institutional Review Board. Patients were enrolled at MedStar Georgetown University Hospital and the University of Utah. Both men and women were eligible for participation; however, all enrolled patients were women. All participants provided informed consent.

### Study Treatment

All participants received a four-week induction of talazoparib 1 mg PO daily on days 1-28. Beginning with cycle 2, patients received a combination of talazoparib 1 mg PO daily on days 1-28 plus avelumab 800 mg IV on days 1 and 15 of each 28-day cycle until disease progression or unacceptable toxicity. Research biopsies were obtained at three predefined time points: (1) baseline (BS), prior to initiating talazoparib treatment; (2) after completion of talazoparib monotherapy and before initiation of combination therapy (BX2); and (3) after treatment with the combination regimen (BX3). Collected specimens were used for transcriptomic and spatial biological analyses **(Fig. 1A)**.

### Assessments

The primary objective of the study was to evaluate the safety and tolerability of talazoparib in combination with avelumab. Participants were monitored for safety every two weeks on day 1 and 15 of each cycle for the first four cycles. If no grade 2 or higher toxicities occurred, participants transition to safety evaluations every four weeks, coinciding with day 1 of each cycle.

Secondary endpoints included ORR, PFS, OS, CBR and duration of response (DoR). Restaging exams were performed every eight weeks based on calendar time rather than cycle number. For participants who experienced a complete response, partial response, or stable disease confirmed on two consecutive restaging assessments, imaging intervals were extended to every 12 weeks. Participants remained on study as long as there was no evidence of disease progression and study therapy was adequately tolerated. Interim analyses for PFS and OS were prespecified at 12 and 24 months. Participants demonstrating ongoing clinical benefit beyond two years of therapy were transitioned to talazoparib monotherapy.

### Statistics

The patient was treated as the independent biological unit throughout this study. Repeated biopsies, cells, spatial neighborhoods, fields of view, and regions of interest were not treated as mutually independent patient-level replicates; where individual cells, regions, or repeated timepoints were analyzed, non-independence was addressed using mixed-effects models with patient identity as a random effect, or by aggregating to patient– or sample-level summaries prior to testing.

### Statistical analysis for clinical and survival data

Descriptive statistics were used to summarize patients’ baseline demographics and adverse events. Kaplan-Meier methodology was used to analyze time-to-event endpoints (PFS and OS), and Log-rank test was used for group comparison. ORR and CBR were calculated as the proportion of patients achieving the specified response category among all treated patients within each cohort; patients without a formally evaluable best response were classified as non-responders for these calculations. Among BRCA-WT patients, two had a best response that was not evaluable (one not evaluable, one too early for assessment at the time of data cutoff). p-value <0.05 is considered statistically significant. SAS software (SAS Ins., Gary, NC) Version 9.4 was used for statistical analysis.

### Statistical analysis of correlative multi-omic data

Unless otherwise specified, comparisons of continuous outcomes (gene expression, protein expression, cell-type frequencies, marker-positive fractions) used linear mixed-effects models with biopsy timepoint as a fixed effect and patient identity as a random effect, fit separately within each BRCA group (see platform-specific Methods above); contrasts (BX2−BS, BX3−BS) were derived using estimated marginal means. All LMER-based P values are two-sided.

Associations between continuous variables and PFS, and between PD-1 expression and local cell density (**Fig. 4H, 4K**; **Supplementary Fig. 6A-C, E, G**), were assessed using simple linear regression, fit separately within each subgroup and timepoint (**Fig. 4H**; **Supplementary Fig. 6A-C, E, G**) or on cells pooled across all patients and samples without a patient-level random effect (**Fig. 4K**). P values (two-sided, from the regression slope) and R² are reported per facet; no correction was applied across facets.

Associations between marker expression and local neighboring-cell density at the single-cell level (**Fig. 4K, 6I, 6J, 6N**; **Supplementary Fig. 10A, G-I**) were assessed using simple linear regression on cells pooled across all patients and samples, for illustrative purposes. These P values reflect cell-level, not patient-level, sample size and are not corrected for within-patient correlation. The hierarchically appropriate, patient/cluster-level treatment of these same relationships — including between-patient heterogeneity in the density-expression relationship — is provided by the Bayesian mixed-effects model in **Fig. 4L** and the derived patient-level slope analyses in **Supplementary Fig. 6F**.

Pairwise associations between continuous variables are reported as Spearman’s ρ or Pearson’s r (**Fig. 3F**, **4J**, **6L**), without an accompanying significance test. For analyses where a formal hypothesis test was performed, correction for multiple comparisons (Benjamini–Hochberg) was applied within specific, pre-defined test families and is stated explicitly in the corresponding figure legend; where a figure legend does not state a correction method, P values are nominal and reflect the exploratory nature of these correlative analyses, uncorrected for multiple testing.

Normality of model residuals was assessed separately for each platform’s model family (IO360 gene-level, DSP Protein, and CyCIF cell-type/marker-level) by pooling standardized residuals (each fit’s residuals divided by its own residual SD) across all fits within that platform. Because pooled sample sizes were large, Shapiro-Wilk significance was evaluated across 20 independent random subsamples (n=1,000) per platform, reporting the test statistic (W) rather than the p-value, which is highly sensitive to sample size. IO360 residuals showed no meaningful departure from normality (W median=0.997). DSP Protein and CyCIF residuals showed modest departures — a slight heavy-tailed (S-shaped) Q-Q pattern for DSP Protein (W median=0.994) and a more pronounced leptokurtic pattern for CyCIF (W median=0.957) — neither judged to materially bias the mixed-model estimates or their standard errors, given the robustness of LMER point estimates to moderate non-normality at these sample sizes.

Effect sizes are reported as the model coefficient (log2 fold-change or slope) for LMER and linear regression analyses, the odds ratio for Fisher’s exact tests, the normalized enrichment score (NES) for GSEA, and Spearman’s ρ or Pearson’s r for correlation analyses, with exact per-comparison values provided in the Source Data files accompanying each figure.

The statistical test, sidedness, and correction applied to each main and supplementary figure panel are listed in **Supplementary Table 3**.

### Data acquisition and experimental procedures

Tumor biopsies were profiled using complementary platforms capturing transcriptomic, protein-level, and spatial single-cell features.

Bulk transcriptomic profiling was performed using the NanoString PanCancer IO360 panel supplemented with 55 DNA damage–related genes. Tumor-enriched regions were microdissected from FFPE sections following pathologist annotation. Spatial profiling was performed using NanoString GeoMx Digital Spatial Profiling (DSP) Protein Assays, and cyclic immunofluorescence (CyCIF). For DSP protein analysis, protein expression was quantified separately within pan-cytokeratin–positive (tumor) and pan-cytokeratin–negative (stromal) compartments.

For CyCIF, FFPE sections underwent iterative rounds of immunofluorescence staining and imaging using a RareCyte CyteFinder HT microscope (20×/0.75 NA). Antibodies used for CyCIF targeted epithelial, immune, and cell-state markers to enable multiplexed spatial profiling. The panel included pan-cytokeratin (panCK; Cell Signaling Technology, Cat# 4545), CD3 (Abcam, Cat# ab16669), CD4 (Abcam, Cat# ab133616), CD8 (Dako, Cat# M7103), CD68 (Dako, Cat# M0814), and CD163 (Novus Biologicals, Cat# NBP2-39067) to define tumor and immune compartments. Immune checkpoint and cytotoxic markers included PD-1 (BioLegend, Cat# 329906), PD-L1 (Cell Signaling Technology, Cat# 13684), and Granzyme B (Thermo Fisher Scientific, Cat# MA1-80734). Cell-state and signaling markers included phospho-TBK1 (Ser172; Cell Signaling Technology, Cat# 5483), γH2AX (Ser139; MilliporeSigma, Cat# 05-636), cleaved caspase-3 (Asp175; Cell Signaling Technology, Cat# 9661), and phospho-ERK1/2 (Thr202/Tyr204; Cell Signaling Technology, Cat# 4370). Nuclear staining was performed using Hoechst 33342 (Thermo Fisher Scientific, Cat# H3570). All antibodies were validated for compatibility with iterative CyCIF staining and stripping cycles, and signal specificity was confirmed by expected spatial localization and consistency across samples.

All datasets were annotated with patient identifiers, cohort information, and biopsy time points prior to integrative analysis.

### Quality control and sample inclusion criteria

All correlative analyses were performed on samples that met predefined, platform-specific quality control (QC) criteria. QC was assessed independently for each assay, and failure of a sample in one modality did not preclude its inclusion in other analyses. Patient inclusion for correlative analyses was therefore determined at the level of individual assays rather than requiring complete cross-platform overlap. The number of samples analyzed per platform, genotype, and timepoint is summarized in **Supplementary Table 4**. Each biopsy specimen was profiled once per platform (no technical replicate averaging); longitudinal biopsies from the same patient were treated as repeated measures via patient-level random effects in all mixed-effects models (see below).

### Patient-level inclusion and exclusion for correlative analyses

Patients were included in correlative analyses if at least one biopsy specimen (baseline or on-treatment) passed QC in at least one profiling modality (IO360, DSP Protein, or CyCIF). All samples meeting assay-specific QC criteria were included irrespective of clinical response, biopsy time point, or availability of matched samples at other time points.

Two patients were excluded from correlative analyses based on biological considerations that could confound interpretation of treatment-induced effects. Patient #1017 had received prior treatment with abatacept, a CTLA4 agonist, which can induce durable immune modulation and interfere with interpretation of subsequent immune responses. Patient #1004, assigned to the BRCA–WT cohort was the only patient in the entire 24-patient trial to undergo the protocol-specified fourth research biopsy (BX4), performed following disease progression after prior clinical benefit — an atypical clinical course for the BRCA-WT cohort that prompted genomic review, which identified a frameshift mutation in *BLM*, a DNA damage repair gene mechanistically implicated in BRCA-like sensitivity to PARP inhibition.^48, 49^ Given this finding, patient #1004 was excluded from the primary genotype-stratified correlative (multi-omic) analyses to avoid confounding interpretation of BRCA-MUT versus BRCA-WT tumor microenvironment differences; this patient’s clinical outcome remains included in the efficacy analyses reported in **Fig. 1**.

### Assay-specific quality control

#### IO360

Bulk transcriptomic data were generated and QC-filtered using standard NanoString PanCancer IO 360 pipelines. All samples passing manufacturer-defined internal QC metrics were included in downstream analyses.

#### CyCIF

CyCIF analyses included only samples that yielded interpretable signal through up to six staining cycles. Samples with substantial tissue loss, pervasive staining artifacts, or failed segmentation were excluded. Analyses were restricted to markers available in the successfully completed cycles for each sample.

#### GeoMx DSP Protein Assays

DSP protein data were QC-filtered at the AOI level. AOIs failing platform-defined QC metrics were excluded, while samples were retained if at least one AOI passed QC. All qualified AOIs from retained samples were included in downstream analyses.

### Cross-platform concordance and integrated CD8 analysis

Cross-platform concordance was assessed by comparing shared analytes across IO360, DSP protein and CyCIF datasets. To quantify CD8 T cell dynamics, IO360 CD8A expression and CyCIF CD8 T cell abundance were standardized and integrated using mixed-effects models, with DSP protein serving as an orthogonal validation.

### IO360 bulk transcriptomic analysis

#### Transcriptomic data processing

Bulk transcriptomic analyses were performed using normalized NanoString PanCancer IO360 expression data generated using the standard NanoString pipeline. Expression matrices were integrated with curated clinical annotations, including biopsy time point, BRCA1/2 mutation status, and cohort information, to enable longitudinal and cross-sectional analyses.

#### Gene-level and pathway-level analyses

Gene-level analyses were performed using linear mixed-effects models to assess associations with biopsy time point and BRCA status. Time point was modeled as a fixed effect, and patient identity was included as a random intercept to account for repeated measurements. Both paired (within-patient longitudinal) and unpaired (cohort-level) comparisons were accommodated within the same modeling framework. Because no direct cross-genotype (BRCA-MUT vs. BRCA-WT) comparison was performed at the gene level — every contrast was within-genotype (BX2 or BX3 versus baseline, within one BRCA group at a time) — models were fit separately within each BRCA group rather than including genotype as a covariate in a single combined model, avoiding an unnecessary assumption of shared residual and random-effect variance across genotypes that may differ.

For visualization and downstream ranking, model-derived statistics were converted to signed association scores defined by the direction and magnitude of the effect (sign of coefficient × – log10 P value). These signed statistics were used to generate gene-level summary matrices across contrasts.

Pathway-level analyses were performed using curated gene sets, including NanoString IO360-defined pathways, Hallmark gene sets, and custom gene sets related to DNA damage response and innate immune signaling (including cGAS–STING). Gene set enrichment analysis was performed using a preranked framework based on signed association statistics. In parallel, pathway-level activity scores were also analyzed directly using mixed-effects models, ensuring consistency between gene-level and pathway-level inference.

### DSP spatial protein analysis (GeoMx)

#### Definition of tumor and stromal compartments

Spatial protein expression was measured using NanoString GeoMx Digital Spatial Profiling (DSP). Regions of interest (ROIs) were selected on FFPE sections, and areas of illumination (AOIs) were defined using pan-cytokeratin staining. PanCK-positive AOIs were designated as tumor, and panCK-negative AOIs as stroma, enabling compartment-resolved protein quantification within each sample.

#### AOI-level modeling, aggregation, and cohort analysis

Protein expression values (IgG-normalized and log2-transformed) were analyzed primarily at the AOI level. Linear mixed-effects models were used with patient identity as a random effect and AOI treated as a repeated measurement nested within patient. This framework enabled estimation of longitudinal effects (BX2 and BX3 versus baseline) and genotype-associated differences while accounting for intra-sample heterogeneity. BRCA status, tissue segment (tumor or stroma), and time point were combined into a single interaction factor within each model, enabling direct comparisons across genotype and segment as well as across time points, with patient identity included as a random effect to account for repeated within-patient sampling.

Given the high concordance of protein expression between tumor and stromal AOIs within individual samples (**Supplementary Fig.4A,B**), per-sample protein summaries were generated by averaging AOI-level signals across compartments for analyses requiring a single representative value. AOI-level data were retained for analyses explicitly modeling spatial compartmentalization.

### CyCIF spatial single-cell analysis

#### Image preprocessing, segmentation, and data generation

Multiplexed CyCIF images were processed using the MCMICRO pipeline, including image registration, illumination correction, and cell segmentation. Single-cell marker intensities, spatial coordinates, and metadata were extracted and assembled into analysis-ready objects using the CycifAnalyzeR framework.

#### Marker signal processing and gating

For each sample, a high-quality region of interest (ROI) was manually defined using OMERO to exclude regions affected by tissue damage, staining artifacts, or imaging defects. Marker intensities were inspected and thresholded on a per-marker basis to distinguish signal from background. Rather than performing explicit global normalization, batch– and sample-level variability were accounted for within downstream statistical models.

#### Cell type calling and lineage definition

Cell types were assigned using marker-based classification combined with spatial context. Major lineages included tumor, immune, and stromal cell types (**Supplementary Fig. 2C**).

Tumor cells were classified as panCK-high or panCK-low populations. panCK-high tumor cells were defined by strong epithelial marker expression. In contrast, panCK-low tumor cells were identified primarily based on spatial context, defined as cells residing within high-density tumor regions and exhibiting spatial continuity with panCK-high tumor cells. This approach enabled retention of tumor cells with reduced epithelial marker expression while excluding isolated non-tumor cells.

Marker expression was used as a secondary criterion to exclude cells with strong lineage-specific signals inconsistent with tumor identity. Thus, panCK-low tumor classification reflected spatial embedding within tumor-core structures rather than marker expression alone.

Immune cell types were defined based on canonical marker expression, including CD8+ T cells, CD4+ T cells, and CD3+CD4−CD8− T cells (T_other), as well as macrophage populations defined by CD68, CD163, and double-positive states. Stromal cells were defined as non-epithelial, non-immune cells lacking lineage-defining markers.

This combined marker– and spatially informed framework enabled consistent annotation of cell types across samples while preserving biologically relevant heterogeneity in tumor and immune populations.

#### Sample-level summaries and statistical modeling

Single-cell annotations were aggregated to generate per-sample metrics, including cell-type frequencies, marker-positive fractions, and tumor–stroma composition. Longitudinal and cohort-level comparisons were performed using linear mixed-effects models with patient as a random effect. BRCA status and time point were combined into a single interaction factor within each model, enabling direct within– and across-genotype comparisons at each time point, while patient identity was included as a random effect to account for repeated measurements across time points.

#### Cellular neighborhood construction and spatial metrics

Cellular neighborhoods were defined using a fixed-radius approach, identifying all cells within 20 μm of each index cell. Neighborhood-level features included local cell-type composition and average marker expression.

Neighborhood-level composition metrics were computed for each cell by quantifying the relative frequency of neighboring cell types within a fixed-radius (20 μm) neighborhood. These metrics captured local cellular mixing and were used to define microenvironmental states and to model associations between cell-type composition and marker expression.

#### Dimensionality reduction and clustering of macrophages and cellular neighborhoods

Unsupervised analyses were restricted to macrophage populations and cellular neighborhood–level features.

For macrophage analysis, cells were pooled across samples with subsampling (maximum 2,000 cells per sample) to prevent dominance by high-cell-yield samples. Uniform Manifold Approximation and Projection (UMAP) was used for dimensionality reduction based on lineage and state-marker expression, followed by Leiden clustering to define 10 macrophage states.

For neighborhood-level analysis, all neighborhoods were pooled across samples and embedded using UMAP based on cell type composition and marker features. Leiden clustering identified 17 TME clusters. Cluster number was selected based on stability and interpretability and held constant across analyses.

#### Spatial regression analysis

Local density values for each cell type and marker used in these analyses were computed by smoothing raw local neighborhood counts using k-nearest-neighbor averaging (k = 20 neighbors) in the relevant feature space, reducing noise from finite local cell counts prior to downstream regression modeling.

Associations between marker expression in index cells and local neighborhood features were assessed using hierarchical modeling frameworks. Linear mixed-effects models were used for most analyses, with patient identity modeled as a random effect.

Group-level estimates and post-hoc pairwise comparisons from these mixed-effects models were obtained using estimated marginal means (R package emmeans), which compute model-adjusted group means and contrasts while accounting for the fitted fixed– and random-effect structure.

For analyses of PD-1 regulation, two Bayesian hierarchical multilevel models (brms, Student-t likelihood) were fit. The primary model (**Fig. 4L**) included PD-1 expression in T cells as the outcome, with center cell type (CD8+, CD4+, or T_other) entered as a fixed interaction with local CD8+, CD4+, and T_other T cell density, a random intercept for patient, and a random intercept and slope for each density term by BRCA status × timepoint group (uncorrelated random effects). This model was fit to 6,270 cells from 14 patients. A complementary model, fit separately for each center cell type using the same predictors and random-effects structure, was used to obtain partially pooled group-level slopes (**Supplementary Fig. 6E**) and, for the CD8+ T cell subtype, partial residuals for each density term (**Supplementary Fig. 6F**). These models were fit separately for each T cell subtype using cells from the same 14 patients (CD4+ T cells: n = 2,238; CD8+ T cells: n = 2,673; T_other T cells: n = 1,359). Default weakly informative priors (brms defaults) were used for both models. Each model was fit with 4 chains of 4,000 iterations (2,000 warmup); adapt_delta was raised to 0.95 for the primary model. Convergence was assessed via R-hat and the number of divergent transitions: primary model, R-hat = 1.003, 4 of 8,000 post-warmup draws divergent; CD4+ T cell subtype model, R-hat = 1.002, 21 divergent transitions; CD8+ T cell subtype model, R-hat = 1.003, 25 divergent transitions; T_other T cell subtype model, R-hat = 1.003, 16 divergent transitions.

For **Fig. 4L**, point estimates and 95% credible intervals for each cell-type × density combination were obtained from the fixed-effect coefficients, and pairwise comparisons between neighboring-density coefficients within each cell type were assessed from the posterior draws (twice the smaller one-sided tail probability, as a two-sided Bayesian analog of a P value). For **Supplementary Fig. 6E**, group-level (BRCA status × timepoint) coefficients from the complementary model were obtained as the fixed effect plus each group’s partially pooled deviation, and pairwise comparisons between neighboring-density group-level coefficients were assessed using paired Wilcoxon signed-rank tests (paired by BRCA status × timepoint group). For **Supplementary Fig. 6F**, as a representative example, partial residuals were computed for the CD8+ T cell subtype by subtracting, from PD-1 expression, the fitted contribution of the two non-focal densities using that group’s own coefficients from the **Supplementary Fig. 6E** model, then plotted against the focal density within each BRCA status × timepoint group.

By contrast, **Fig. 4K, 6I, 6J, 6N**, and **Supplementary Fig. 10A, G, H, and I** display individual cells pooled across all samples and patients for visualization, without correcting for patient-level clustering; these panels are illustrative rather than the primary statistical evidence, which instead comes from the hierarchical models above that explicitly account for nested cell-, neighborhood-, timepoint-, and patient-level structure. Degrees of freedom and P values for linear mixed-effects models throughout this study were computed using the Satterthwaite approximation, the default method for lmerTest-fitted models.

#### Integrated transcriptomic and spatial single-cell analysis

Integration analyses focused on IO360 transcriptomic data and CyCIF-derived single-cell features to maximize sample coverage. After confirming concordance between platforms for CD8-related measurements, per-sample CD8A expression (IO360) and CyCIF-derived CD8+ T cell frequency were each standardized (z-scored) and combined into a single integrated CD8 signal per sample using a mixed-effects model with sample identity as a fixed effect and platform (technology) as a random effect. This approach enabled unified analysis across samples lacking complete cross-platform overlap while preserving platform-specific signal structure. Longitudinal and genotype-stratified analyses were then performed on this integrated signal using the same linear mixed-effects framework applied throughout the correlative analyses (biopsy timepoint × BRCA status as a fixed effect, patient identity as a random effect; see Statistical analysis of correlative multi-omic data).

#### Murine Brca1-Deficient TNBC Model and Flow Cytometric Analysis of Tumor-Infiltrating Immune Cells

Brca1-deficient TNBC cells (904), derived from a tumor arising in the *Ade-CMV-Cre Brca1^fl/fl^ p53^fl/fl^* genetically engineered mouse model and their isogenic Brca1 addback counterparts (904 Brca1+) were orthotopically implanted into the fourth mammary fat pad of 6–8-week-old female FVB/n mice. Tumor-bearing mice were treated daily with either vehicle control (DMSO) or olaparib (50 mg/kg) for 14 consecutive days. All animal studies were approved by the Institutional Animal Care and Use Committee (IACUC) at Brigham and Women’s Hospital, under protocol # 2020N000142 and performed in accordance with institutional guidelines and regulations.

At the end of treatment, tumors were harvested and enzymatically dissociated using the MACS Tumor Dissociation Kit (Miltenyi Biotec, 130-096-730) according to the manufacturer’s instructions. Cell suspensions were washed, subjected to red blood cell lysis, and counted. Cells were resuspended in PBS and plated into 96-well U-bottom plates for downstream staining.

Prior to antibody staining, cells were blocked with anti-mouse Fcγ receptor II/III antibody (CD16/CD32; Affymetrix, 14-0161-85) and stained with a fixable live/dead dye (Zombie NIR; BioLegend) for 20 min at room temperature in the dark. Cells were then stained with fluorophore-conjugated antibodies against CD45 (eFluor450, clone 30-F11; Invitrogen, 48-0451-82), CD3 (BV570, clone 17A2; BioLegend, 100225), and CD8a (BV510, clone 53-6.7; BioLegend, 100751) for 1 h at 4 °C protected from light.

Following staining, cells were washed with FACS buffer (PBS containing 0.5% BSA and 2 mM EDTA), fixed using a commercial fixation buffer (Invitrogen, 00-8222-49) for 20 min at room temperature, and resuspended in FACS buffer for analysis. Data were acquired on an Aurora spectral flow cytometer (Cytek), and flow cytometry data were analyzed using OMIQ software (Dotmatics). The gating strategy for flow cytometric identification of tumor-infiltrating CD8+ T cells is shown in **Supplementary Fig. 11**.

#### LLM assistance disclosure

During preparation of this manuscript, the authors used ChatGPT (OpenAI) and Claude (Anthropic) to assist with writing and debugging analysis code, as well as to assist with drafting and editing portions of the manuscript text, including administrative submission materials (e.g., Reporting Summary). The analytical approach and study design were determined by the authors; the tools were not used to draw scientific conclusions. All code and text were reviewed and validated by the authors, who take full responsibility for their accuracy and content.

## Data Availability

Bulk transcriptomic (NanoString IO360) and spatial protein (NanoString GeoMx DSP) data are deposited in GEO under accession numbers GSE339263 and GSE339236; single-cell spatial (CyCIF) data are deposited at Zenodo under https://doi.org/10.5281/zenodo.21815064. These datasets are under embargo and will be made publicly available upon publication in a peer-reviewed journal. For all statistical tests, only significance thresholds (P or FDR values) are indicated in figure panels and legends; the corresponding point estimates (e.g., regression coefficients, correlation coefficients, group means) and their associated uncertainty (e.g., standard errors, confidence intervals) are provided in the Source Data files accompanying each figure. Individual-level clinical data are not publicly available due to patient privacy but are available from the corresponding authors upon reasonable request, subject to institutional data use agreements.

## Code Availability

Analysis code is available on GitHub at https://github.com/GuerrieroLab/TALAVE_project, tagged ‘biorxiv-preprint’ for the version cited in this manuscript.

## Acknowledgements/funding

We thank the patients and their families for their participation in this study. We thank the Dana-Farber Cancer Institute Molecular Pathology Core Laboratory for GeoMx DSP assay execution and data acquisition, the Brigham and Women’s Hospital Center for Advanced Molecular Diagnostics Research Core for NanoString IO360 processing and Digital Analyzer support, and NanoString Technologies (Bruker Spatial Biology) for PanCancer IO360 transcriptomic profiling and GeoMx Digital Spatial Profiling analysis support. Talazoparib and avelumab were provided by Pfizer, as part of an alliance between Pfizer and the healthcare business of Merck KGaA, Darmstadt, Germany.

This work was supported by the Dana-Farber/Harvard Cancer Center SPORE in Breast Cancer (NIH grant P50 CA168504), the Susan G. Komen Foundation (CCR18547597), NCI Cancer Systems Biology Center of Excellence Grant (U54-CA225088), Terri Brodeur Breast Cancer Foundation, The Friends of Dana-Farber, The Harvard Ludwig Center, NIH grant R01/R37 CA269499 (to J.L.G.), NIH grant R01 CA307291 (to J.L.G., F.L., and G.I.S.), and in part by NCI Cancer Center Support Grants to the Georgetown Lombardi Comprehensive Cancer Center (P30 CA051008) and the Dana-Farber/Harvard Cancer Center (P30 CA006516).

## Author Contributions

Conceptualization: F.L., C.I., X.G., H.W., J.S.B., G.I.S., J.L.G

Methodology: F.L., K.S., C.I., X.G., E.T.R., A.N., K.F.Z., M.G.T., S.J.S., H.W., J.S.B., J.L.G.

Investigation: F.L., C.I., X.G., A.N., E.T.R., P.R.P., K.F.Z., M.G.T., N.S., S.J.S., H.W., G.I.S.

Formal analysis: X.G., H.W., K.S., K.F.Z., M.G.T., F.L., C.I., J.S.B., G.I.S., J.L.G.

Data curation: K.S., E.T.R., S.J.S.

Software: K.S., X.G., H.W.

Resources: K.S., J.S.B., J.L.G.

Project administration: F.L., C.I., X.G., H.W., G.I.S., J.L.G.

Funding acquisition: F.L., G.I.S., J.L.G.

Supervision: F.L., C.I., G.I.S., J.L.G.

Writing – original draft: F.L., K.S., G.I.S., J.L.G.

Writing – review & editing: F.L., K.S., C.I., X.G., E.T.R., A.N., C.M., M.W., J.M.C., P.R.P., A.L.H., K.F.Z., M.G.T., N.S., L.M.S., J.S., S.J.S., H.W., J.S.B., G.I.S., J.L.G.

## Competing Interests

F.L. provides consulting/advisory board services for Pfizer, AstraZeneca, Daiichi Sankyo and receives grant/research support (institution) from: AstraZeneca, Gilead, Incyte, Zentalis, Ideaya and Merck.

K.S. serves on the Scientific Advisory Board of FELIQS Corporation.

E.T.R. is currently employed by Merck & Co., Inc, Rahway, NJ, USA, a shareholder of Merck & Co., Inc, Rahway, NJ, USA, and an inventor on a patent pending entitled “Deep Learning Enabled Slide Image Analysis for Prediction of Treatment Response” assigned to Merck & Co., Inc, Rahway, NJ, USA.

M.W. provides consulting/advisory board service for Lilly, AstraZeneca, Gilead, Merck, Novartis, Jazz, Genetech Relay Therapeutics. Research funding from Incrediwear (paid to Institute).

A.L.H. provides consulting/advisory board services for AstraZeneca, Daiichi Sankyo, and Genentech/Roche.

M.G.T. is currently employed by Moderna, Inc. Moderna had no role in this study or its publication.

J.S.B. is a member of the SAB of Frontier Medicines and Dialectic Therapeutics.

G.I.S. has research funding from Merck KGaA/EMD-Serono, Artios, Lilly and Pfizer. He has served on advisory boards for Merck KGaA/EMD-Serono, Circle Pharmaceuticals, Concarlo Therapeutics, Schrodinger, FoRx Therapeutics and MycRx. He holds patents entitled, “Dosage regimen for sapacitabine and seliciclib,” and “Compositions and Methods for Predicting Response and Resistance to CDK4/6 inhibition.”

J.L.G. provides consulting services for Antana Bio LLC, Array BioPharma/Pfizer, AstraZeneca, BD Biosciences, BigHat, Duke Street Bio, Genentech, GlaxoSmithKline, Incyte, iTeos, Laverock, LTZ, OncoOne, Synkine, and Voro; and receives Grant/Research support from: Array BioPharma/Pfizer, Duke Bioscience, Eli Lilly, GlaxoSmithKline, Merck.

P.R.P. receives grant/research support (to institution) from: Jazz and ALX Oncology. She has served on advisory boards for Pfizer, Personalized Cancer Therapy (Perthera), Sirtex, Heron, Puma, BOLT, AbbVie.

