## Supplementary information for "TALAVE in Breast Cancer: *BRCA1/2* Mutation-Dependent Immune Remodeling after PARP Inhibition with Limited Checkpoint Engagement"

The following is provided as separate an Excel spreadsheet file:

**Supplementary Data 1.** Custom gene panel and pathway annotations for NanoString IO360 transcriptomic profiling.

**Supplementary Figure 1. PFS and OS outcomes highlight differential therapeutic activity in BRCA-MUT and BRCA-WT cohorts**

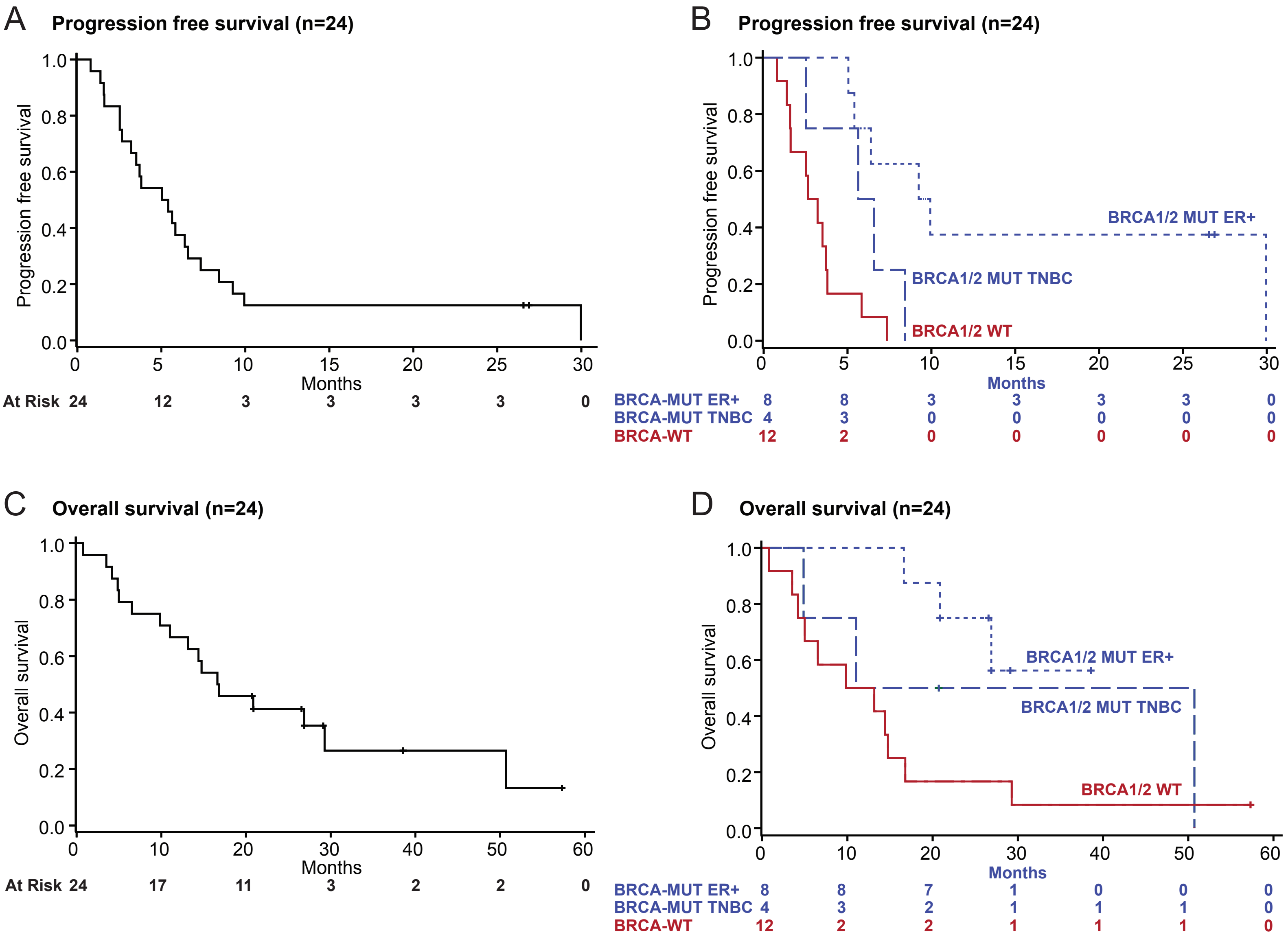

**Supplementary Figure 1. PFS and OS outcomes highlight differential therapeutic activity in BRCA-MUT and BRCA-WT cohorts.**

(A,B) Kaplan-Meier curve showing the median PFS for (A) the overall study population and (B) the population stratified by BRCA status and disease subtypes.

(C,D) Kaplan-Meier curve showing the median OS for (C) the overall study population and (D) the population stratified by BRCA status and disease subtypes.

P values are from two-sided log-rank tests comparing PFS or OS between the indicated groups (see Methods).

### Supplementary Figure 2. Therapy induces structural remodeling of BRCA-MUT but not BRCA-WT tumors

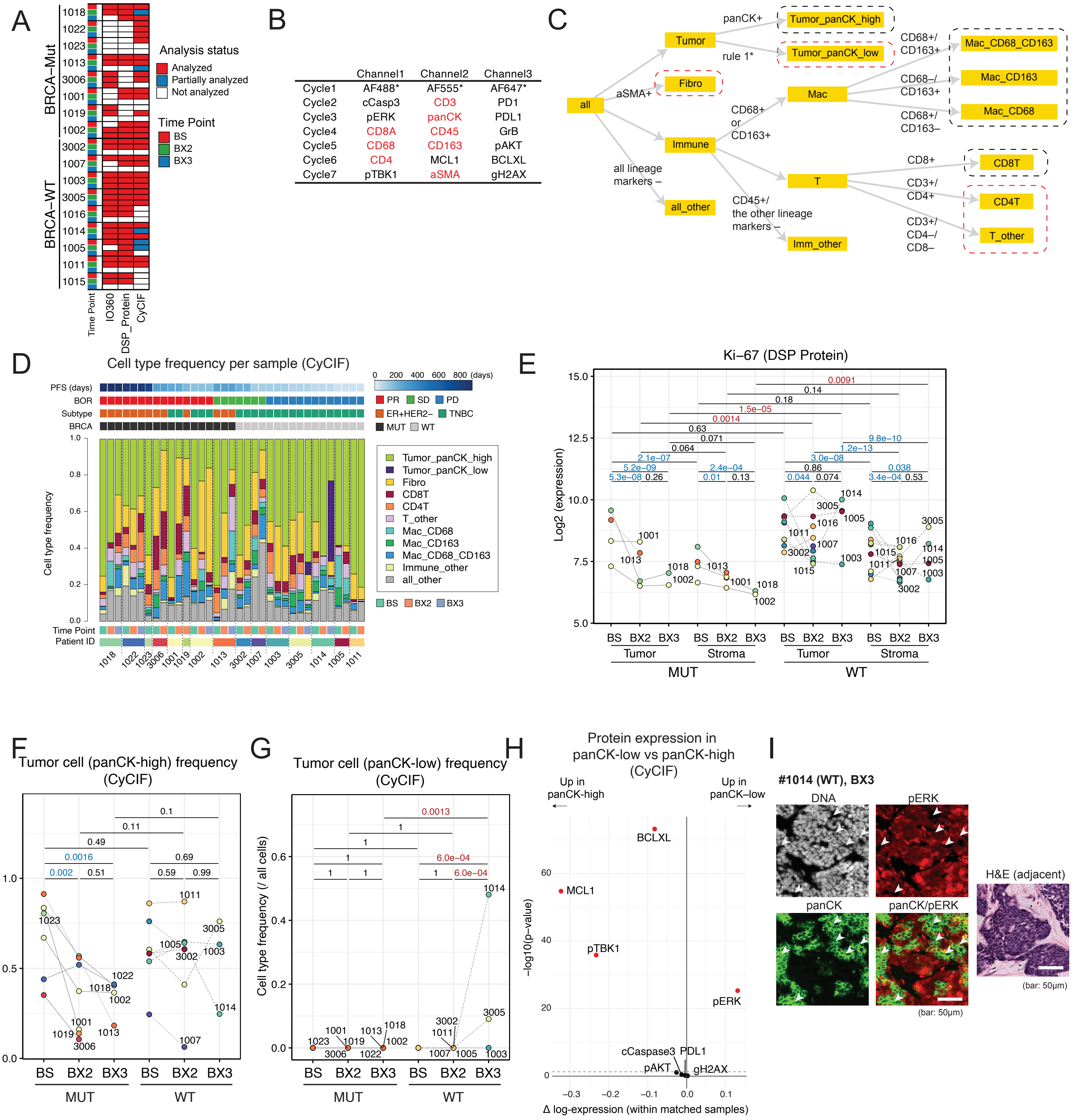

**Supplementary Figure 2. PARP inhibition depletes tumor cells and disrupts tumor architecture in BRCA-MUT but not BRCA-WT tumors.**

(A) Sample distribution across three technologies. In IO360 and DSP Protein Assays, red and white indicate analyzed and excluded samples. In CyCIF, red and blue denote data available through cycles 7 and 5, respectively. Patients are ordered by BRCA status followed by PFS.

(B) List of antibodies measured in each staining cycle. Markers highlighted in red indicate those used for cell type classification; those in black denote cell state markers. Cycle 1 included three autofluorescence channels for background correction, without antibody staining.luded three autofluorescence channels at different wavelengths without antibody staining for background correction.

(C) Hierarchical classification of cell types based on positive marker expression, applied incrementally from left to right. Cell types in black boxes require lineage marker expression through cycle 7 and are therefore unavailable in samples limited to cycles 1–5; cell types in red boxes require marker expression available already by cycle 5 and are classified in all samples.

(D) Stacked bar plots of cell type frequencies for individual samples, stratified by BRCA status, best overall response, and PFS.

(E) Mean Ki-67 expression per patient measured by DSP Protein Assays, stratified by BRCA status, ROI (tumor vs. stroma), and time point.

(F) Frequency of panCK-high tumor cells as a proportion of all cells.

(G) Frequency of panCK-low tumor cells as a proportion of all cells.

(H) Volcano plot showing differential expression of cell state markers between panCK–high and panCK–low tumor cells in matched samples (n = 2). Red points indicate markers with significant differences.

(I) Representative images of panCK-low tumor cells in CyCIF, showing panCK (green), phospho-ERK (red), and DNA (white), with H&E staining of an adjacent section included for reference.

P values in E, F, and G are from two-sided linear mixed-effects models with patient as a random effect. H uses the same LMER framework, but with sample (not patient) as the random effect, given the cell-state comparison is made within each of only 2 matched samples; points denote nominal $P < 0.05$, uncorrected. See Methods.

**Supplementary Figure 3. Talazoparib suppresses MMEJ pathways in BRCA-MUT but not BRCA-WT tumors**

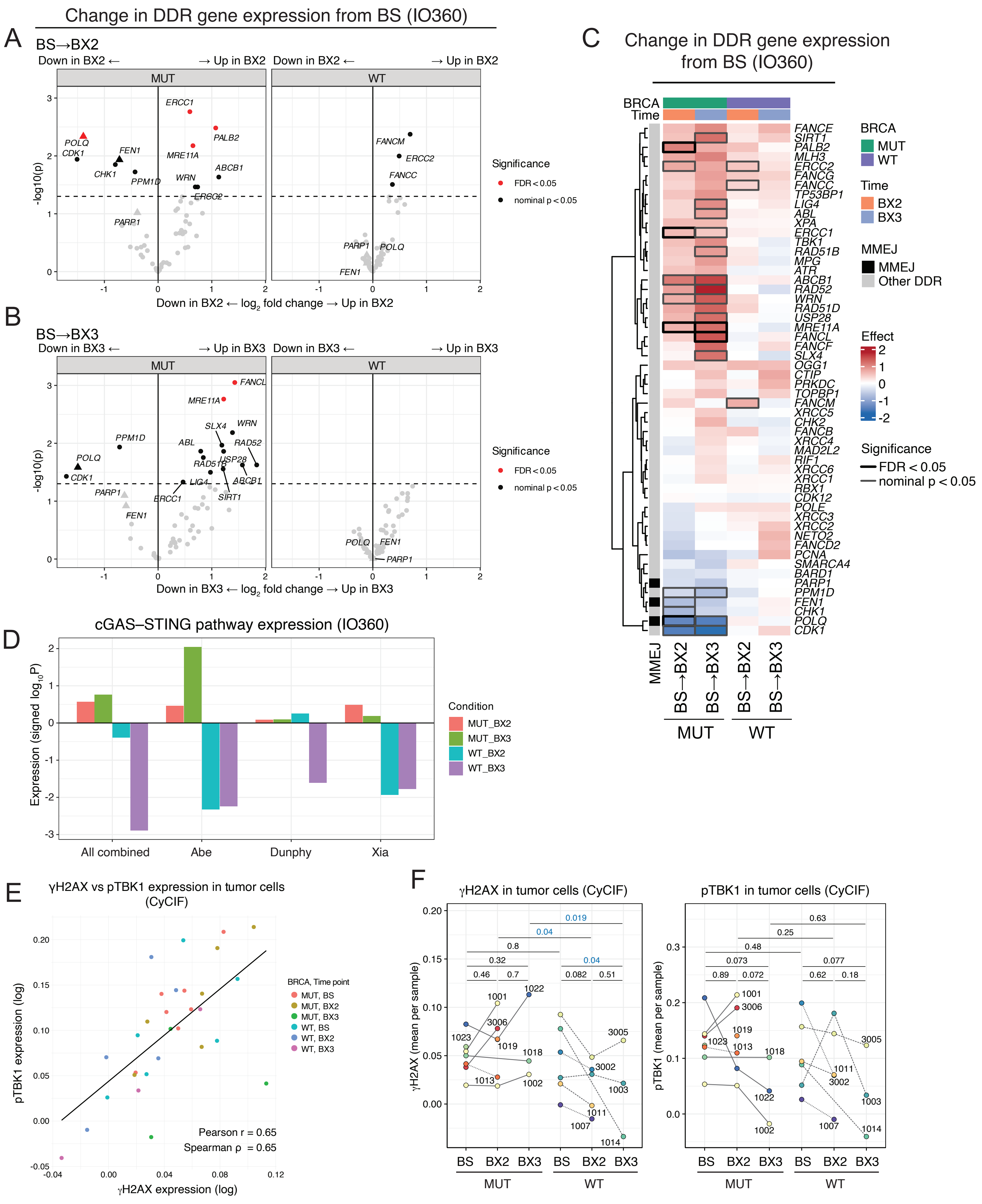

**Supplementary Figure 3. Talazoparib suppresses MMEJ pathways in BRCA-MUT but not BRCA-WT tumors.**

(A–B) Volcano plots showing differential expression of DNA damage response (DDR) genes from baseline (BS) to BX2 (A) and BX3 (B) in BRCA-MUT and BRCA-WT tumors (IO360 RNA profiling). MMEJ-associated genes (PARP1, FEN1, POLQ) are marked by shape (filled triangle).

(C) Heatmap of DDR gene expression across patients, shown as change from baseline. Genes involved in MMEJ are indicated in black on the left.

(D) Gene set enrichment of cGAS–STING signatures from individual references.

(E) Scatterplot of mean tumor-cell γH2AX vs. pTBK1 expression per patient sample (Pearson r and Spearman ρ shown); colors indicate BRCA status and sampling time point.

(F) Temporal changes in mean tumor-cell γH2AX and pTBK1 expression across biopsy timepoints, stratified by BRCA status.

In A and B, the y-axis shows nominal $-\log10(P)$; point color reflects a three-tier significance scheme (red: $FDR < 0.05$; black: nominal $P < 0.05$ only; grey: not significant). In C, box border denotes $FDR < 0.05$ (black) or nominal $P < 0.05$ only (grey). In D, bar height reflects signed $-\log10(FDR)$, as computed by fgsea. In F, P values are from two-sided linear mixed-effects models with patient as a random effect. See Methods.

**Supplementary Figure 4. UMAP of all cells profiled by CyCIF imaging**

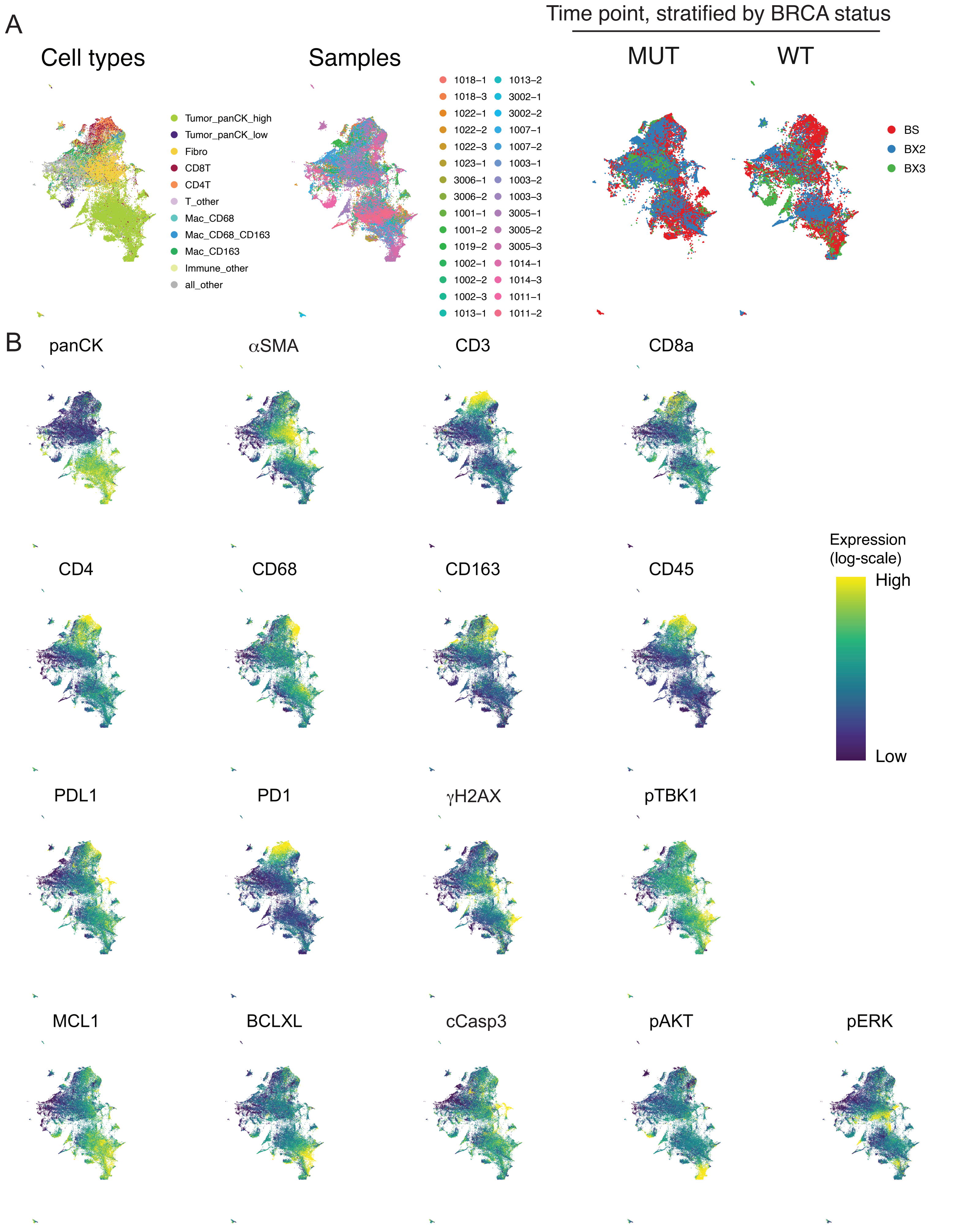

**Supplementary Figure 4. UMAP of all cells profiled by CyCIF imaging.**
(A) UMAPs of all cells across all samples, colored separately by cell type, BRCA status, treatment time point, sample ID, and experimental batch (one UMAP per variable).
(B) UMAPs colored by expression of selected lineage and cell state markers (one UMAP per marker).

#### Supplementary Figure 5. Broad immune-related protein marker expression was attenuated in BRCA-WT tumors upon PARP inhibition

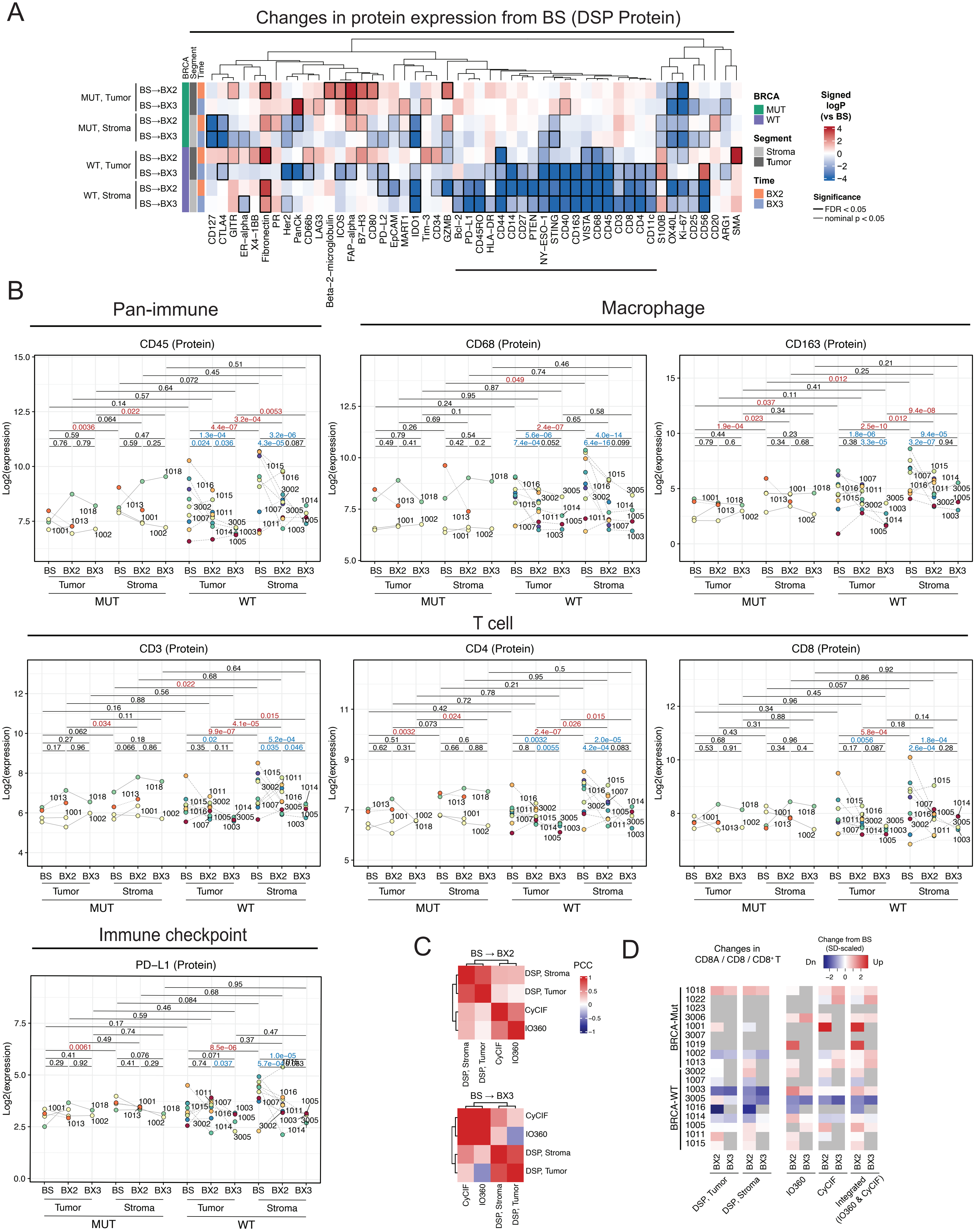

**Supplementary Figure 5. Broad immune-related protein marker expression was attenuated in BRCA-WT tumors upon PARP inhibition.**

(A) Heatmap of DSP protein markers showing treatment-induced changes from baseline to BX2 and BX3, stratified by tumor–stroma compartment and BRCA status. Colors indicate signed $\log_{10}$(nominal P) values from linear mixed-effects models. Box border denotes FDR < 0.05 (black) or nominal P < 0.05 only (grey). Proteins marked by line segments indicate treatment-associated downregulation in BRCA-WT tumors and correspond to multiple immune-related markers.

(B) Seven lineage markers (pan-immune, macrophage, and T cell, Immune checkpoint) showing reduced expression following treatment in BRCA-WT tumors, with no significant changes observed in BRCA-MUT tumors.

(C) Heatmaps showing concordance of gene and protein expression changes from BS to BX2 (top) and BX3 (bottom) across platforms and patients.

(D) Longitudinal changes in CD8-associated measurements across DSP Protein Assays, IO360, CyCIF, and integrated IO360–CyCIF analysis per patient.

P values in A and B are from two-sided linear mixed-effects models with patient as a random effect. In A, color reflects nominal P and box border denotes FDR < 0.05 (black) or nominal P < 0.05 only (grey); in B, P values are nominal, not corrected across the 7 markers shown (individual-protein display, as in Fig. 3E/Supplementary Fig. E2E). C reports Pearson correlation coefficients, no formal test. See Methods.

### Supplementary Figure 6. BRCA-dependent remodeling of the TME links PD-1 expression in T cells to CD4+ T cell density and PFS

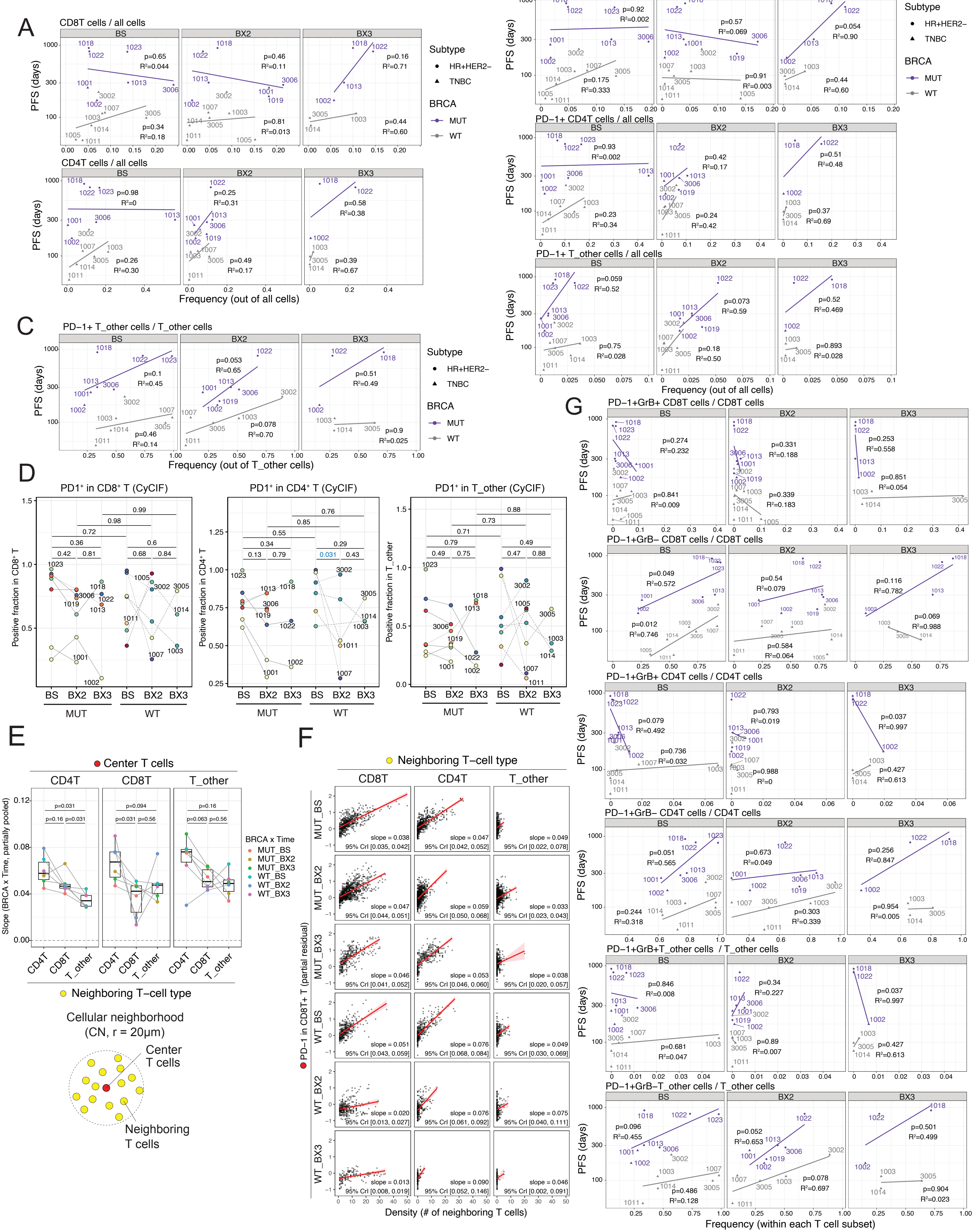

**Supplementary Figure 6. BRCA-dependent remodeling of the TME links PD-1 expression in T cells to CD4+ T cell density and PFS.**

(A) Relationship between the frequency of CD8+ and CD4+ T cells out of all cells and PFS.

(B) Relationship between the fractions of PD-1+ cells for each T cell subtype out of all cells and PFS.

(C) Relationship between the fraction of PD-1+ cells out of all T_other cells and PFS.

(D) Fraction of PD1+ cells in CD8T, CD4T, and T_other per patient, stratified by BRCA groups and time points.

(E) Group-level regression slopes for PD-1 expression versus CD8+, CD4+, or T_other T cell density, modeled hierarchically with partial pooling across BRCA status × timepoint subgroups; brackets denote paired comparisons across neighboring T cell types within each center cell type.

(F) As a representative example, association between PD-1 expression in CD8+ T cells and local CD8+, CD4+, or T_other T cell density, adjusted for the other two densities (partial residual), stratified by BRCA status × timepoint subgroup.

(G) Relationship between the fractions of PD-1+GrB+ and PD-1+GrB- cells within each T cell subtype and PFS.

P values in D are from two-sided linear mixed-effects models with patient as a random effect. P values in A, B, C, and G are from two-sided linear regression. E reports paired Wilcoxon signed-rank tests comparing density-term slopes within each T cell subtype (paired by BRCA status × timepoint group). F shows partial residuals from the hierarchical model underlying E, for the CD8+ T cell subtype only, as a representative example; no formal test performed. See Methods.

#### Supplementary Figure 7. Changes in cell type frequencies during treatment measured by CyCIF

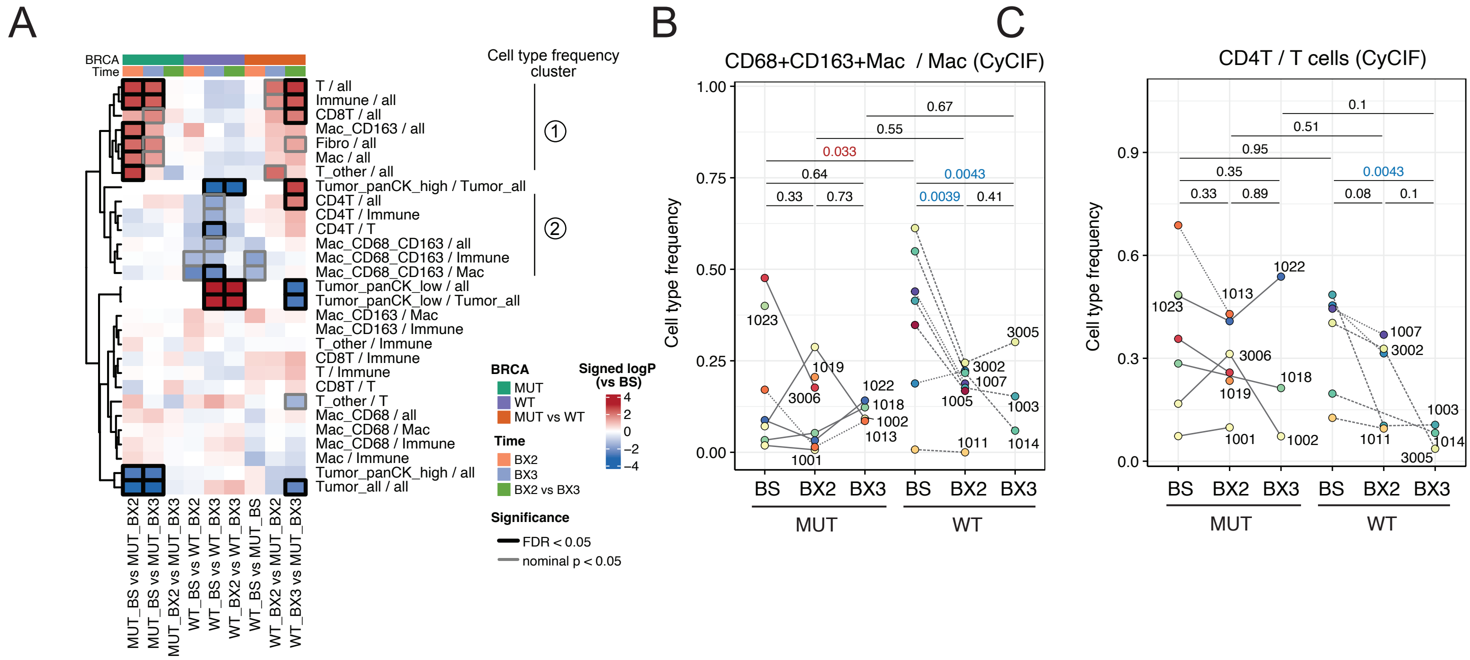

**Supplementary Figure 7. Changes in cell type frequencies during treatment measured by CyCIF.** (A) Heatmap of cell type frequency changes across treatment and BRCA status. Each column represents a comparison: BRCA-MUT (left), BRCA-WT (middle), and BRCA-WT vs. BRCA-MUT (right). Colors show signed $\log_{10}$ p values from linear mixed-effects models (red: higher in latter condition; blue: higher in former of each contrast). The two cell type clusters identified in Fig. 5A (Cluster 1/Cluster 2) are grouped together here. (B,C) Frequency of CD68+/CD163+ double-positive macrophages within the macrophage compartment (B); frequency of CD4+ T cells within the T cell compartment (C). P values are BH-adjusted for multiple comparisons in A; see Methods.

**Supplementary Figure 8. Macrophage subtypes show distinct spatial distribution, tumor-stroma ratios, and temporal dynamics**

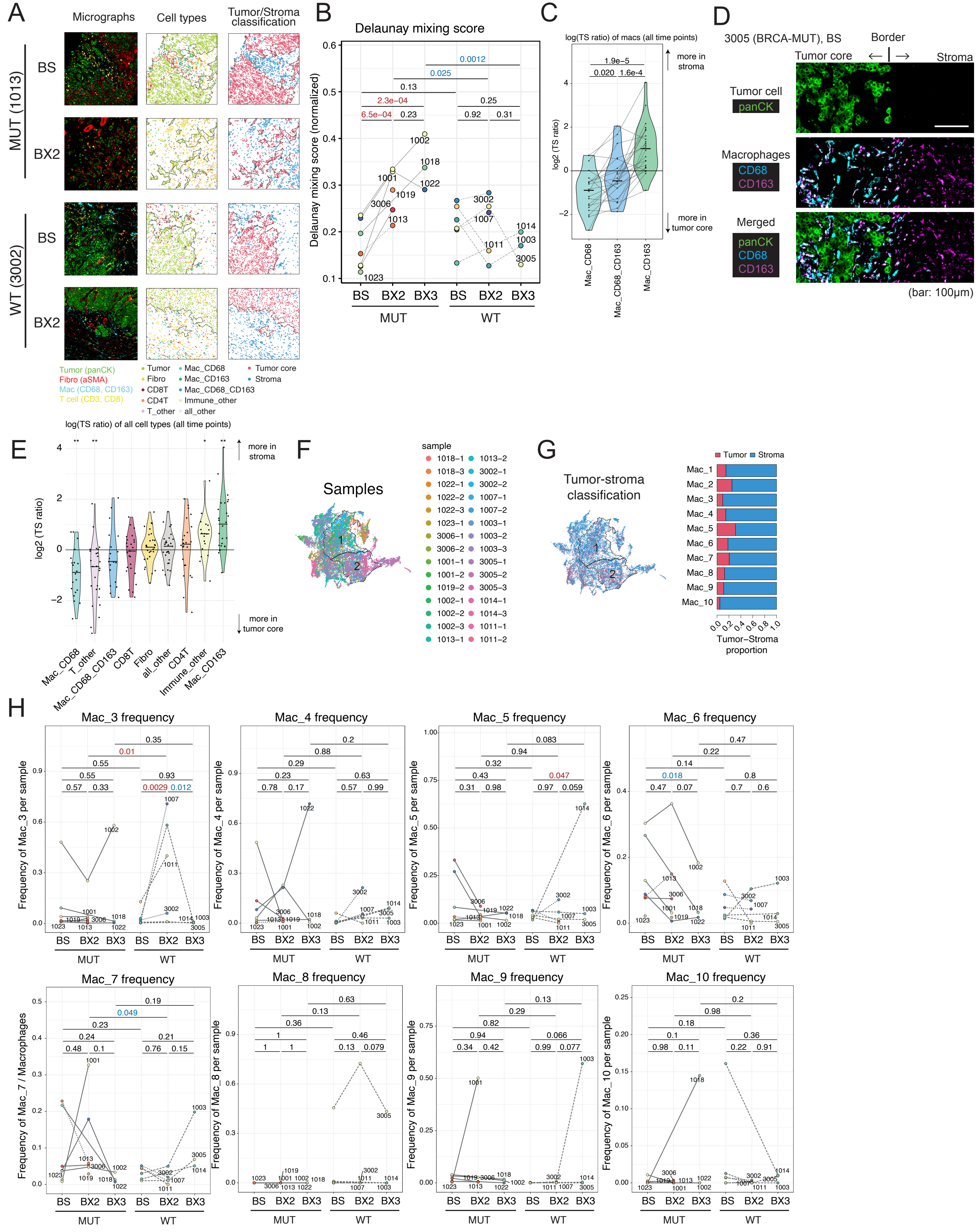

**Supplementary Figure 8. Macrophage subtypes show distinct spatial distribution, tumor-stroma ratios, and temporal dynamics.**

(A) Tumor–stroma classification based on CN cell-type frequencies. For each row (same region across columns): left, raw micrograph; middle, cells colored by computationally assigned cell type; right, tumor vs. stroma classification predicted by a logistic classifier trained on CN-level cell-type composition. Top rows: BRCA-MUT (1013); bottom rows: BRCA-WT (3002); each shown at BS and BX2.

(B) Delaunay mixing score (normalized), quantifying tumor–stroma intermingling, over time by BRCA status. The score increased over time in BRCA-MUT tumors, indicating progressive disintegration of tumor–stroma boundaries, while remaining low and stable in BRCA-WT tumors.

(C) Tumor-to-stroma ratio (TS ratio) for macrophage subtypes (CD68+, CD163+, CD68+/CD163+ double-positive). Horizontal line within each violin denotes the median.

(D) Representative micrographs (sample 3005, BRCA-MUT, BS) showing tumor core, border, and stroma regions, with panCK (tumor), CD68, and CD163 (macrophages) and merged channels. Dashed lines delineate the tumor-core/stroma border. (Scale bar: 100 μm.)

(E) TS ratio of all measured cell types. Horizontal line within each violin denotes the median.

(F) UMAP of macrophages colored by sample, with Mac_1 and Mac_2 cluster boundaries indicated by dashed lines.

(G) Predicted spatial localization of macrophages in tumor core versus stroma. Left: UMAP of macrophages colored by predicted location (tumor core, blue; stroma, red). Right: bar plot showing the proportion of macrophages in tumor core versus stroma.

(H) Relative frequency of Mac_3 through Mac_10 macrophages among all macrophages, stratified by BRCA status and time point.

P values in B are from two-sided linear mixed-effects models with patient as a random effect. P values in C are from two-sided paired Wilcoxon signed-rank tests comparing TS ratio between macrophage subtypes within the same sample, uncorrected. Asterisks in E denote two-sided one-sample Wilcoxon signed-rank tests (H0: median log2(TS ratio) = 0) within each cell type, uncorrected; $^*P < 0.05$, $^{**}P < 0.01$. See Methods.

### Supplementary Figure 9. Characterization of TME cluster features

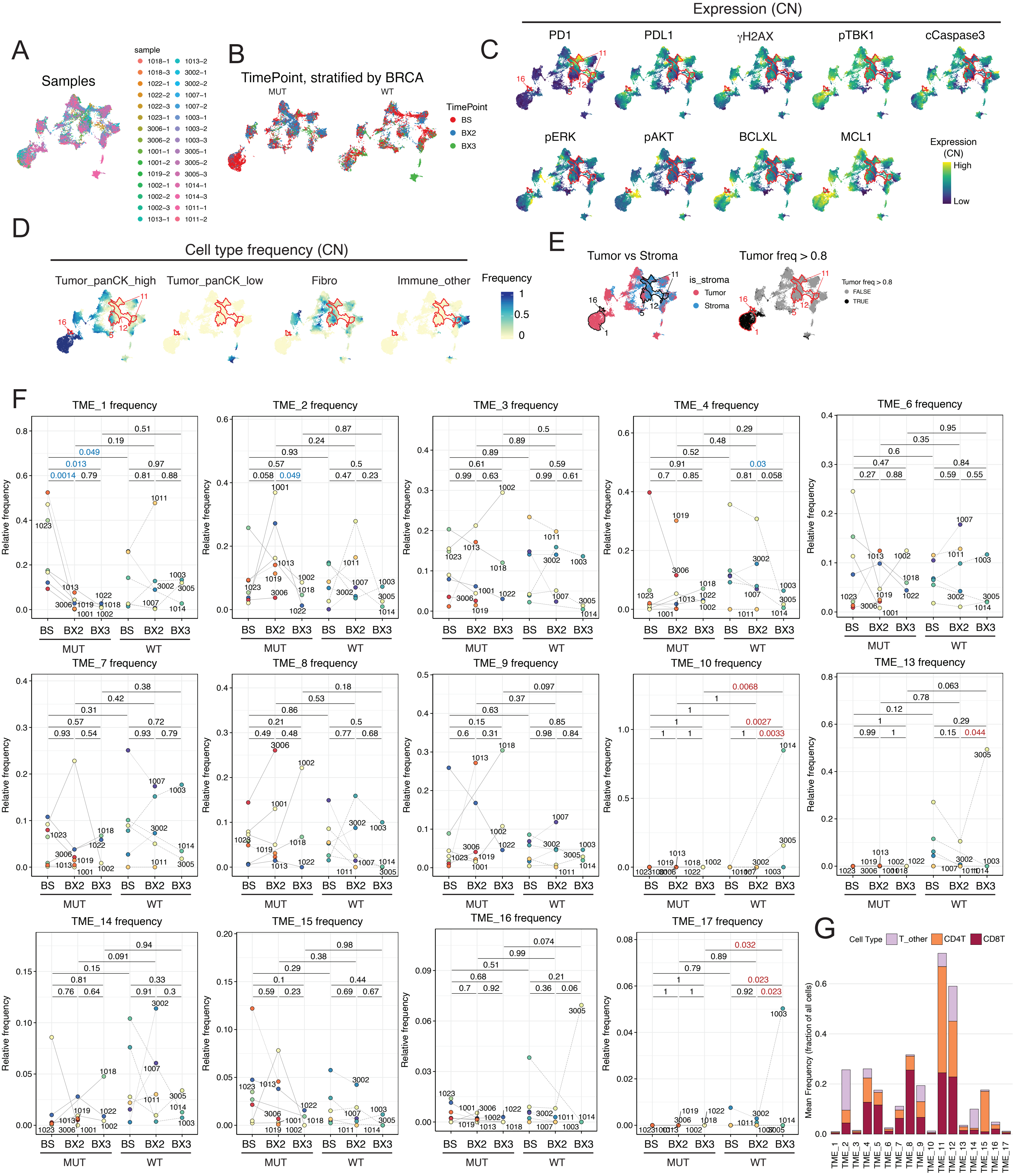

**Supplementary Figure 9. Characterization of TME cluster features.**

(A–B) UMAP of CNs colored by (A) sample and (B) time point, shown separately for BRCA-MUT and BRCA-WT tumors.

(C–D) UMAPs colored by (C) selected protein expression (one UMAP per protein) and (D) frequencies of neighboring cells from each cell type (one UMAP per cell type).

(E) UMAPs colored by focal-cell designation (tumor vs. stroma; left) and by CN tumor purity (CNs with >80% tumor cells; right).

(F) Frequency of each TME cluster stratified by BRCA status and time point.

(G) Mean fraction of T cells per TME cluster. T cells are color-coded by subtype.

P values in F are from two-sided linear mixed-effects models with patient as a random effect. See Methods.

### Supplementary Figure 10. CN Clustering reveals immunologically distinct PD-L1+ microenvironments

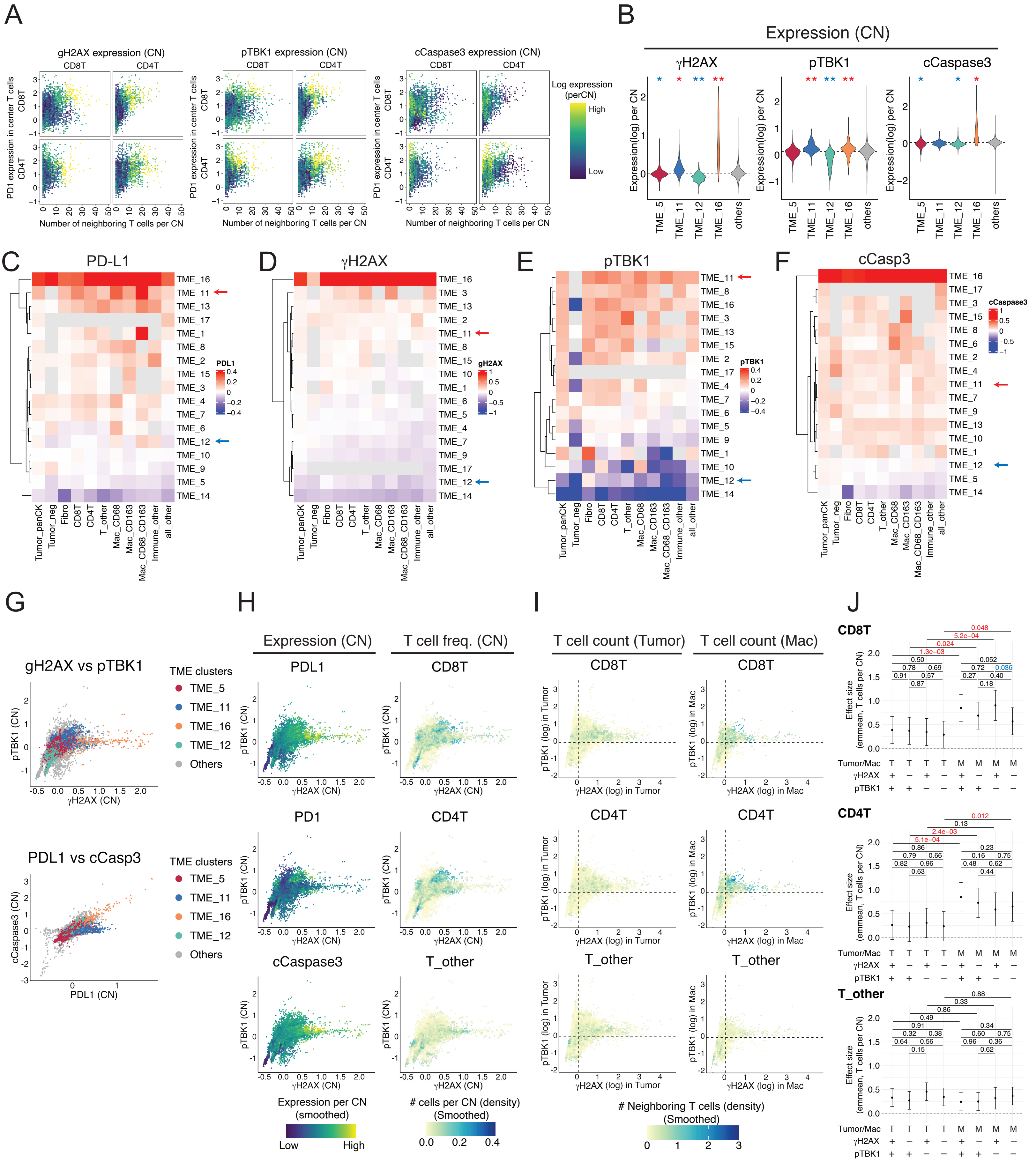

**Supplementary Figure 10. CN Clustering reveals immunologically distinct PD-L1+ microenvironments**.

(A) Cell state marker expression at the CN level overlaid on the relationship between PD-1 expression in T cells and the number of neighboring T cells within 20-μm CNs (CD4+ and CD8+ T cells only): γH2AX (left), phospho-TBK1 (middle), and cCasp3 (right).

(B) Expression of selected proteins in CNs belonging to four selected TME clusters (TME_5, TME_11, TME_12, TME_16) and all remaining CNs combined.

(C-F) Mean cell state marker expression per cell type and TME cluster (grey indicates missing data): (C) PD-L1, (D) γH2AX, (E) pTBK1, (F) cCasp3.

(G) CN-level expression of co-expressed protein pairs: pTBK1/γH2AX, and PD-L1/cCasp3. Four key TME clusters are indicated.

(H) CN-level expression of pTBK1 and γH2AX, colored by protein expression (left) and by the number of neighboring T cells (right).

(I) Single-cell expression of pTBK1 and γH2AX in tumor cells (left) and macrophages (right). Each point represents an individual cell and is colored by the number of neighboring T cells (CD8+, CD4+, or T_other). Dashed lines delineate marker-positive and marker-negative quadrants.

(J) Linear mixed-effects model coefficients summarizing differences in neighboring T cell density across the pTBK1/γH2AX-defined regions shown in (I).

P values are BH-adjusted for multiple comparisons in B; P values in A, G, H, and I are from cell-level regression, shown for illustration only (see Methods and Fig. 4L for the patient-level treatment); pairwise contrasts in J are nominal P values, uncorrected. See Methods.

**Supplementary Figure 11. Gating strategy for flow cytometric identification of tumor-infiltrating CD8+ T cells.**

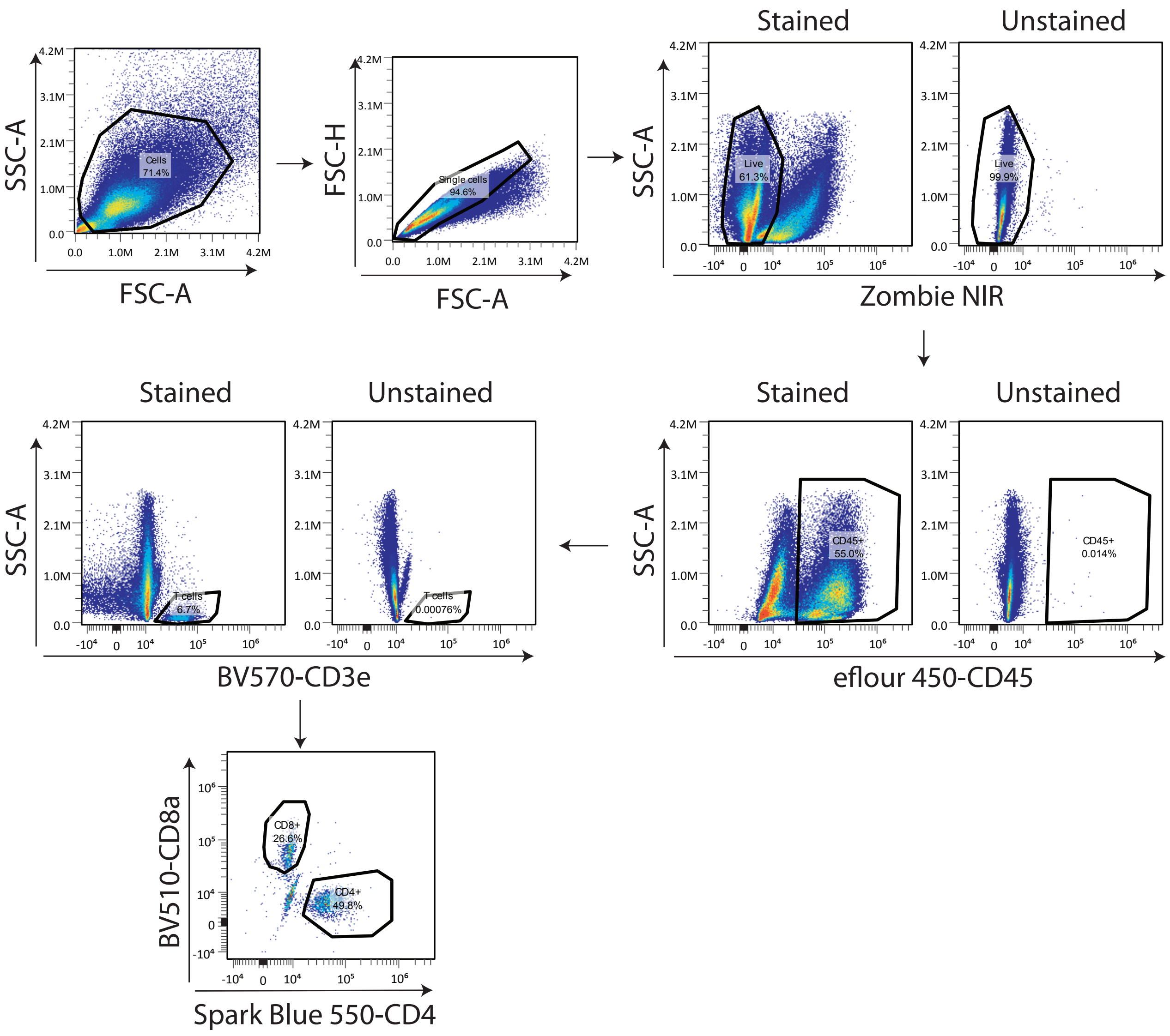

**Supplementary Figure 11. Gating strategy for flow cytometric identification of tumor-infiltrating CD8+ T cells.**

Representative gating strategy used to identify tumor-infiltrating CD8+ T cells in mouse by flow cytometry. Cells were sequentially gated on FSC-A vs. SSC-A (cells), FSC-A vs. FSC-H (single cells), viability (Zombie NIR−, live cells), and CD45 (immune cells), followed by CD3e (T cells) and CD4/CD8a co-staining to identify CD4+ and CD8+ T cell subsets.

**Supplemental Table 1. Baseline patient characteristics.**

| Characteristic | Cohort 1 (n = 12) | Cohort 2 (n = 12) | Overall (N = 24) |
|---|---|---|---|
| Median age, years (range) | 45.5 (27 – 66) | 56.5 (37 – 78) | 50 (27 – 78) |
| Race | | | |
| White | 8 (66.7%) | 10 (83.3%) | 18 (75%) |
| Black or African American | 3 (25%) | 1 (8.3%) | 4 (16.7%) |
| Unknown | 1 (8.3%) | 1 (8.3%) | 2 (8.3%) |
| Ethnicity | | | |
| Non-Hispanic | 11 (91.7%) | 11 (91.7%) | 22 (91.7%) |
| Unknown | 1 (8.3%) | 1 (8.3%) | 2 (8.3%) |
| Prior lines of therapy for metastatic disease | | | |
| 0 | 5 (41.7%) | 4 (33.3%) | 9 (37.5%) |
| 1 | 5 (41.7%) | 3 (25%) | 3 (12.5%) |
| ≥2 | 2 (16.7%) | 5 (41.7%) | 7 (29.2%) |
| Last Known Survival Status | | | |
| Alive | 9 (75%) | 2 (16.7%) | 11 (45.8%) |
| Dead | 3 (25%) | 10 (83.3%) | 13 (54.2%) |
| Progression or Death | | | |
| No | 2 (16.7%) | 0 (0%) | 2 (8.3%) |
| Yes | 10 (83.3%) | 12 (100%) | 22 (91.7%) |

**Supplemental Table 2. Treatment-related adverse events.**

| Adverse Event No. of patients (%) | Cohort 1 (n = 12) | | Cohort 2 (n = 12) | | Overall (N = 24) | |
|---|---|---|---|---|---|---|
| | **All-Grade** | **Grade 3-4** | **All-Grade** | **Grade 3-4** | **All-Grade** | **Grade 3-4** |
| Fatigue | 3 (25%) | 0 (0%) | 6 (50%) | 0 (0%) | 9 (37.5%) | 0 (0%) |
| Nausea | 4 (33.3%) | 0 (0%) | 5 (41.7%) | 0 (0%) | 9 (37.5%) | 0 (0%) |
| Anemia | 5 (41.7%) | 4 (33.3%) | 3 (25%) | 3 (25%) | 8 (33.3%) | 7 (29.2%) |
| Neutropenia | 5 (41.7%) | 2 (16.7%) | 1 (8.3%) | 1 (8.3%) | 6 (25%) | 3 (12.5%) |
| Thrombocytopenia | 2 (16.7%) | 1 (8.3%) | 3 (25%) | 2 (16.7%) | 5 (20.8%) | 3 (12.5%) |
| Rash maculo-papular | 1 (8.3%) | 0 (0%) | 3 (25%) | 0 (0%) | 4 (16.7%) | 0 (0%) |
| Infusion-related reaction | 4 (33.3%) | 0 (0%) | 0 (0%) | 0 (0%) | 4 (16.7%) | 0 (0%) |
| Constipation | 1 (8.3%) | 0 (0%) | 2 (16.7%) | 0 (0%) | 3 (12.5%) | 0 (0%) |
| Fever | 2 (16.7%) | 0 (0%) | 0 (0%) | 0 (0%) | 2 (8.3%) | 0 (0%) |
| Hyperthyroidism | 1 (8.3%) | 0 (0%) | 1 (8.3%) | 0 (0%) | 2 (8.3%) | 0 (0%) |
| Stomach pain | 0 (0%) | 0 (0%) | 2 (16.7%) | 0 (0%) | 2 (8.3%) | 0 (0%) |
| Electrocardiogram QT corrected interval prolonged | 0 (0%) | 0 (0%) | 2 (16.7%) | 0 (0%) | 2 (8.3%) | 0 (0%) |
| Aspartate aminotransferase increased | 1 (8.3%) | 1 (8.3%) | 1 (8.3%) | 0 (0%) | 2 (8.3%) | 1 (4.2%) |
| Leukopenia | 2 (16.7%) | 1 (8.3%) | 0 (0%) | 0 (0%) | 2 (8.3%) | 1 (4.2%) |
| Dyspnea | 0 (0%) | 0 (0%) | 2 (16.7%) | 1 (8.3%) | 2 (8.3%) | 1 (4.2%) |
| Alopecia | 2 (16.7%) | 0 (0%) | 0 (0%) | 0 (0%) | 2 (8.3%) | 0 (0%) |
| Thrombotic thrombocytopenic purpura | 1 (8.3%) | 1 (8.3%) | 0 (0%) | 0 (0%) | 1 (4.2%) | 1 (4.2%) |
| Febrile neutropenia | 1 (8.3%) | 1 (8.3%) | 0 (0%) | 0 (0%) | 1 (4.2%) | 1 (4.2%) |
| Sinus tachycardia | 0 (0%) | 0 (0%) | 1 (8.3%) | 0 (0%) | 1 (4.2%) | 0 (0%) |
| Ear and labyrinth disorders – Other, specify | 0 (0%) | 0 (0%) | 1 (8.3%) | 0 (0%) | 1 (4.2%) | 0 (0%) |
| Abdominal distention | 1 (8.3%) | 0 (0%) | 0 (0%) | 0 (0%) | 1 (4.2%) | 0 (0%) |
| Diarrhea | 1 (8.3%) | 0 (0%) | 0 (0%) | 0 (0%) | 1 (4.2%) | 0 (0%) |
| Gastrointestinal disorders – Other, specify | 1 (8.3%) | 0 (0%) | 0 (0%) | 0 (0%) | 1 (4.2%) | 0 (0%) |
| Gastroparesis | 0 (0%) | 0 (0%) | 1 (8.3%) | 0 (0%) | 1 (4.2%) | 0 (0%) |
| Vomiting | 0 (0%) | 0 (0%) | 1 (8.3%) | 0 (0%) | 1 (4.2%) | 0 (0%) |
| Chills | 1 (8.3%) | 0 (0%) | 0 (0%) | 0 (0%) | 1 (4.2%) | 0 (0%) |
| General disorders and administration site conditions – Other, specify | 0 (0%) | 0 (0%) | 1 (8.3%) | 0 (0%) | 1 (4.2%) | 0 (0%) |
| Portal vein thrombosis | 1 (8.3%) | 0 (0%) | 0 (0%) | 0 (0%) | 1 (4.2%) | 0 (0%) |
| Bruising | 0 (0%) | 0 (0%) | 1 (8.3%) | 0 (0%) | 1 (4.2%) | 0 (0%) |
| Alkaline phosphatase increased | 1 (8.3%) | 0 (0%) | 0 (0%) | 0 (0%) | 1 (4.2%) | 0 (0%) |
| Anorexia | 1 (8.3%) | 0 (0%) | 0 (0%) | 0 (0%) | 1 (4.2%) | 0 (0%) |
| Bone pain | 1 (8.3%) | 0 (0%) | 0 (0%) | 0 (0%) | 1 (4.2%) | 0 (0%) |
| Muscle weakness lower limb | 0 (0%) | 0 (0%) | 1 (8.3%) | 0 (0%) | 1 (4.2%) | 0 (0%) |
| Headache | 1 (8.3%) | 0 (0%) | 0 (0%) | 0 (0%) | 1 (4.2%) | 0 (0%) |
| Pruritis | 0 (0%) | 0 (0%) | 1 (8.3%) | 0 (0%) | 1 (4.2%) | 0 (0%) |
| Hot flashes | 1 (8.3%) | 0 (0%) | 0 (0%) | 0 (0%) | 1 (4.2%) | 0 (0%) |
| Lymphedema | 0 (0%) | 0 (0%) | 1 (8.3%) | 0 (0%) | 1 (4.2%) | 0 (0%) |

**Supplementary Table 3. Statistical test, sidedness, and correction applied to each main and supplementary figure panel.**
Each panel appears in exactly one row below, tagged by its test-family Group in column A (equivalent to the former per-family sheet name / Index tab).
Fig.4D is split into -left (IO360 gene expression) and -right (CyCIF frequency) since the two halves use different test families.

| Group | Panel | Comparison | n | Estimate type | Uncertainty type | Statistic type | df | P adj | Sidedness | Correction method |
|---|---|---|---|---|---|---|---|---|---|---|
| Gene_LMER | Fig.3A,B,C; Fig.4B,C; SFig.3A,B, C | Representative row: identical LMER fit per gene, per BRCA group, per contrast (BX2 vs BS, BX3 vs BS). | 8 MUT / 9 WT patients (per-gene n varies with missing data) | LMER coefficient (Δlog2 expression, emmeans contrast) | SE | t | Satterthwaite (emmeans default for lmerTest models) | BH, within each BRCA × contrast family (~805 genes) | two-sided | BH; volcano/heatmap displays use a three-tier nominal+FDR color/box scheme (see legend) |
| Gene_LMER | Fig.2E; Fig.4D-left | Single gene (MKI67; CD8A), same model as above, per-patient trajectory display. | 8 MUT / 9 WT patients | LMER coefficient (Δlog2 expression, emmeans contrast) | SE | t | | n/a (single gene) | two-sided | none |
| GSEA | Fig.3D; Fig.4A; SFig.3D | Representative row: identical preranked fgsea test per pathway (Hallmark, NanoString, custom cGAS-STING/DDR sets). | 8 MUT / 9 WT patients (gene-level ranking input) | NES (fgsea native) | | NES (also serves as the test statistic; permutation-based, no separate t/z) | | BH (padj, fgsea native) | two-sided (permutation) | BH |
| DSP_Protein_LMER | SFig.5A,B | Representative row: identical LMER per protein, 8 of 24 emmeans contrasts kept (within-arm BS/BX2/BX3, MUT-vs-WT same-segment-timepoint). | same AOI × protein matrix for every protein (no missingness) | LMER coefficient (Δlog2 expression, emmeans contrast) | SE | t | Satterthwaite (emmeans default for lmerTest models) | BH, per contrast, pooled across ~52 proteins | two-sided | BH |
| DSP_Protein_LMER | Fig.3E; SFig.2E | Single protein (STING; Ki-67), same model as above, per-patient trajectory display. | same AOI matrix as SFig.5A,B | LMER coefficient (Δlog2 expression, emmeans contrast) | SE | t | Satterthwaite (emmeans default for lmerTest models) | n/a (single protein, raw/uncorrected overlay) | two-sided | none |

| | | | | | | | | | | |
|---|---|---|---|---|---|---|---|---|---|---|
| CyCIF_Sample_LMER | Fig.2B,C (representative of per-marker time-course family) | Illustrative example (Tumor cell frequency, MUT BX2 vs BS); same model/family applies to every other marker/cell-type/density variable shown across this family. | 8 | LMER coefficient | SE | t | | n/a (family of ~3 per marker) | two-sided | none |
| CyCIF_Sample_LMER | Fig.5A; SFig.7A (multi-celltype heatmap family) | Same model as row above, aggregated across cell types into a heatmap. | varies per cell type | LMER coefficient (signed −log10 p) | SE | t | | BH, per column (BRCA × timepoint contrast, across cell-type rows) | two-sided | BH |
| CyCIF_Sample_LMER | SFig.2H | panCK-high vs. panCK-low tumor cells, 2 matched samples; fixed effect fit at single-cell level. | 2 samples at the random-effect level; many cells per sample feed the fixed effect | LMER coefficient (Δexpression) | SE | t | | n/a (family of 8 markers) | two-sided | none |
| CyCIF_Sample_LMER | SFig.8B | Delaunay-based tumor-stroma mixing score (mix_norm), per patient, same 9-contrast structure as the CyCIF frequency family. | 8 | LMER coefficient | SE | t | | n/a (family of ~3) | two-sided | none |
| Integrated_CrossPlatform | Fig.4E; SFig.5D | Integrated CD8 signal (IO360 CD8A + CyCIF CD8T, z-scored, combined), 9 pairwise contrasts. SFig.5D's other 3 sub-panels are single-platform, covered by their own rows above. | 42 samples (patient-timepoint biopsies; excludes patients 1017, 1004) | LMER coefficient (emmeans contrast) | SE | t | | n/a (same policy as CyCIF_Sample_LMER) | two-sided | none |
| Linear_regression | Fig.4H; SFig.6A,B,C,G | Patient-level: PFS (log days) vs. marker fraction/frequency, fit independently per BRCA × timepoint group. | 6-9 patients per group | Slope (lm() coefficient) | SE | t | | n/a (small family, no correction across groups) | two-sided | none |
| Linear_regression | Fig.4K,M; Fig.6I,J,N; SFig.10A,G,H,I | Cell-level: marker expression vs. local neighboring-cell density, cells pooled across all patients/samples, illustration only. | thousands of cells, pooled across ~17 patients (SFig.2A / Supplementary Table 4) | Slope (lm() coefficient) | SE | t | | n/a (not corrected for patient clustering) | two-sided | none |

| | | | | | | | | | | |
|---|---|---|---|---|---|---|---|---|---|---|
| Bayesian | Fig.4L | PD-1 expression (3 T cell centers, modeled jointly) ~ local CD8+/CD4+/T_other density, fixed center interaction, random intercept (patient) and random slope per density term by BRCA×timepoint group. Representative row for all 9 center × density combinations. | 14 patients; 6,270 cells | Posterior mean (fixed-effect slope, brms fixef()) | Posterior SD | | | n/a (Bayesian; posterior comparisons reported per contrast) | two-sided (95% credible interval) | none |
| Bayesian | SFig.6E,F | Complementary model, fit separately per T cell subtype, same predictors/random effects; group-level slopes (SFig.6E) and, for CD8+ subtype only, partial residuals (SFig.6F). | 14 patients; 2,238 / 2,673 / 1,359 cells (CD4+ / CD8+ / T_other subtype models) | Group-level coefficient (fixed effect + partially pooled deviation, brms coef()) | Posterior SD | V (paired Wilcoxon, SFig.6E only) | | n/a (family of 3 comparisons per subtype) | two-sided | none |
| Coefficient_plots | Fig.6O; SFig.10J | LMER emmeans group mean (neighboring T-cell density), Tumor+Mac combined, by PD-L1/cCasp3 (6O) or pTBK1/γH2AX (10J) classification; one panel per T-cell type. | n=25-30 biopsies/group (14 patients), except 1 group in SFig.10J (Mac gH2AX+/pTBK1-): n=20 biopsies/11 patients | emmeans group mean (lmer(mean_tcells ~ group + (1\|Patient.ID))) | 95% CI (emmeans) | t (pairwise contrasts also shown, nominal) | | | two-sided | none |
| Fisher_exact | Fig.5D,I,J | Cell-type enrichment within macrophage TME clusters, 2×2 table per cluster × cell-type pair from aggregate counts. | D: 30 tests (10 clusters x 3 categories); I,J: 110 tests (10 clusters x 11 categories) | Odds ratio | n/a | n/a | | BH, single family across all pairs | one-sided | BH |
| Wilcoxon | Fig.6C | Cell-type frequency and protein/marker expression, in-cluster vs. out-of-cluster, across 17 TME clusters. | Cell-level; in-cluster n=98-5,379 (of 36,521 total), per cluster | n/a | | W (unpaired Wilcoxon) | | BH, per cell type/protein across clusters | one-sided | BH |

| | | | | | | | | | | |
|---|---|---|---|---|---|---|---|---|---|---|
| Wilcoxon | Fig.6F,G,K; SFig.10B | TME cluster (or marker) vs. all remaining clusters pooled, patient-averaged, paired. | up to 17 patients per marker | Median difference | | V | | BH, per marker across 4 target clusters | two-sided | BH |
| Wilcoxon | SFig.8C | Tumor-to-stroma ratio, macrophage subtype pairs, paired, all timepoints pooled. | Paired n=17-20 per pair | n/a | | V | | n/a (family of 3) | two-sided | none |
| Wilcoxon | SFig.8E | Tumor-to-stroma ratio, one-sample test vs. 0, all cell types, all timepoints pooled. | n=18-30 per cell type | n/a | | V | | n/a (family of 9) | two-sided | none |
| Correlation | Fig.4I; Fig.6L; SFig.5C | Pairwise marker correlations, reported descriptively (density threshold, co-expression, or scatter). | n/a | Pearson's r / Spearman's ρ | | n/a (no formal test) | | n/a | n/a | none |
| Correlation | Fig.3F | Per-gene Pearson r (IO360 vs. DSP Protein), tested against zero per dataset-pair category. | n/a | Pearson's r, summarized per dataset-pair category | | t (shown on plot) | | n/a (small family, one test per category) | two-sided | none |

**Supplemental Table 4. Sample and data availability across profiling platforms by patient and timepoint.**

| Patient ID | Timepoint | BRCA1/2 | IO360 | DSP Protein | CyCIF (5 cycles) | CyCIF (7 cycles) |
|---|---|---|---|---|---|---|
| 1018 | BS | MUT | Yes | Yes | Yes | Yes |
| 1018 | BX2 | MUT | Yes | Yes | Yes | No |
| 1018 | BX3 | MUT | No | Yes | Yes | Yes |
| 1022 | BS | MUT | No | No | Yes | Yes |
| 1022 | BX2 | MUT | No | No | Yes | Yes |
| 1022 | BX3 | MUT | No | No | Yes | Yes |
| 1023 | BS | MUT | No | No | Yes | Yes |
| 1023 | BX2 | MUT | No | No | No | No |
| 1023 | BX3 | MUT | No | No | No | No |
| 3006 | BS | MUT | Yes | No | Yes | Yes |
| 3006 | BX2 | MUT | Yes | No | Yes | Yes |
| 3006 | BX3 | MUT | Yes | No | No | No |
| 1001 | BS | MUT | No | Yes | Yes | Yes |
| 1001 | BX2 | MUT | No | Yes | Yes | Yes |
| 1001 | BX3 | MUT | No | No | No | No |
| 3007 | BS | MUT | No | No | No | No |
| 3007 | BX2 | MUT | No | No | No | No |
| 3007 | BX3 | MUT | No | No | No | No |
| 1019 | BS | MUT | Yes | No | No | No |
| 1019 | BX2 | MUT | Yes | No | Yes | Yes |
| 1019 | BX3 | MUT | No | No | No | No |
| 1002 | BS | MUT | No | Yes | Yes | Yes |
| 1002 | BX2 | MUT | Yes | Yes | Yes | Yes |
| 1002 | BX3 | MUT | Yes | Yes | Yes | Yes |
| 1024 | BS | MUT | No | No | No | No |
| 1024 | BX2 | MUT | No | No | No | No |
| 1024 | BX3 | MUT | No | No | No | No |
| 1021 | BS | MUT | No | No | No | No |
| 1021 | BX2 | MUT | No | No | No | No |
| 1021 | BX3 | MUT | No | No | No | No |
| 1013 | BS | MUT | Yes | Yes | Yes | Yes |
| 1013 | BX2 | MUT | Yes | Yes | Yes | Yes |
| 1013 | BX3 | MUT | No | No | Yes | No |
| 3002 | BS | WT | Yes | Yes | Yes | Yes |
| 3002 | BX2 | WT | Yes | Yes | Yes | Yes |
| 3002 | BX3 | WT | No | No | No | No |
| 1007 | BS | WT | Yes | Yes | Yes | Yes |
| 1007 | BX2 | WT | No | Yes | Yes | Yes |
| 1007 | BX3 | WT | No | No | No | No |
| 1003 | BS | WT | Yes | Yes | Yes | Yes |
| 1003 | BX2 | WT | Yes | Yes | Yes | Yes |
| 1003 | BX3 | WT | Yes | Yes | Yes | Yes |
| 3005 | BS | WT | Yes | Yes | Yes | Yes |
| 3005 | BX2 | WT | Yes | Yes | Yes | Yes |
| 3005 | BX3 | WT | Yes | Yes | Yes | Yes |
| 1016 | BS | WT | Yes | Yes | No | No |
| 1016 | BX2 | WT | Yes | Yes | No | No |
| 1016 | BX3 | WT | No | No | No | No |
| 1014 | BS | WT | Yes | Yes | Yes | Yes |
| 1014 | BX2 | WT | Yes | Yes | Yes | No |
| 1014 | BX3 | WT | Yes | Yes | Yes | Yes |
| 1005 | BS | WT | Yes | Yes | Yes | No |
| 1005 | BX2 | WT | No | Yes | Yes | No |
| 1005 | BX3 | WT | Yes | Yes | No | No |
| 1011 | BS | WT | Yes | Yes | Yes | Yes |
| 1011 | BX2 | WT | Yes | Yes | Yes | Yes |
| 1011 | BX3 | WT | No | No | No | No |
| 1015 | BS | WT | Yes | Yes | No | No |
| 1015 | BX2 | WT | Yes | Yes | No | No |
| 1015 | BX3 | WT | No | No | No | No |
| 1009 | BS | WT | No | No | No | No |
| 1009 | BX2 | WT | No | No | No | No |
| 1009 | BX3 | WT | No | No | No | No |
| 3003 | BS | WT | No | No | No | No |
| 3003 | BX2 | WT | No | No | No | No |
| 3003 | BX3 | WT | No | No | No | No |
